# Mutation of Vsx genes in zebrafish highlights the robustness of the retinal specification network

**DOI:** 10.1101/2022.01.20.477122

**Authors:** Joaquín Letelier, Lorena Buono, María Almuedo-Castillo, Jingjing Zang, Sergio González-Díaz, Rocío Polvillo, Estefanía Sanabria-Reinoso, Ruth Diez del Corral, Stephan C. F. Neuhauss, Juan R. Martínez-Morales

## Abstract

Genetic studies in human and mice have established a dual role for *Vsx* genes in retina development: an early function in progenitors’ specification, and a later requirement for bipolar-cells fate determination. Despite their conserved expression patterns, it is currently unclear to which extent *Vsx* functions are also conserved across vertebrates, as mutant models are available only in mammals. To gain insight into *vsx* function in teleosts, we have generated *vsx1* and *vsx2* CRISPR-Cas9 double knockouts (*vsxKO*) in zebrafish. Our electrophysiological and histological analyses indicate severe visual impairment and bipolar cells depletion in *vsxKO* larvae, with retinal precursors being rerouted towards photoreceptor or Müller glia fates. Surprisingly, neural retina is properly specified and maintained in mutant embryos, which do not display microphthalmia. We show that although important *cis*-regulatory remodelling occurs in *vsxKO* retinas during early specification, this has little impact at a transcriptomic level. Our observations point to genetic redundancy as an important mechanism sustaining the integrity of the retinal specification network, and to *Vsx* genes regulatory weight varying substantially among vertebrate species.

**Brief Summary Statement for use in emailed and online tables of content alerts:** The mutation of *vsx* genes in zebrafish confirms a conserved role in bipolar cells specification across vertebrates, but do not interfere with the specification of the neural retina domain. Our data reveal the unexpected robustness of the genetic network sustaining the identity of the neural retina.

## INTRODUCTION

The organogenesis of the vertebrate eye is a complex multistep process entailing the sequential activation of genetic programs responsible for the initial specification of the eye field, the patterning of the eye primordium into sub-domains, and the determination of the different neuronal types. Although we are far from understanding the precise architecture of the gene regulatory networks (GRNs) controlling eye formation, many of their central nodes have been already identified (Buono & Martinez-Morales, 2020; Fuhrmann, 2010; Heavner & Pevny, 2012; Martinez-Morales, 2016). They comprise transcriptional regulators recruited repeatedly for key developmental decisions at different stages of eye formation, and which mutation in humans is often associated to severe ocular malformations: i.e. microphthalmia, anophthalmia, and coloboma. This is the case for SIX3, PAX6, RAX, SOX2, VSX2, or OTX2 (Gregory-Evans et al., 2004; Gregory-Evans et al., 2013).

Among the main regulators, the visual system homeobox transcription factors, Vsx1 and Vsx2, have been shown to control the development of visual circuits in vertebrate and invertebrate species (Burmeister et al., 1996; Erclik et al., 2008; Focareta et al., 2014). *Vsx2*, initially termed as *Chx10*, was the first gene of the family characterized in vertebrates (Liu et al., 1994). *Vsx2/Chx10* shows a conserved expression pattern across vertebrate species, both in the retina (i.e. early in all optic cup precursors, and later in retinal bipolar cells), as well as in hindbrain and spinal cord interneurons (Ferda Percin et al., 2000; Kimura et al., 2013; Liu et al., 1994; Passini et al., 1997). A nonsense mutation in *Vsx2* (Y176stop) turned to be the molecular cause of the phenotype exhibited by the classical mutant mice *ocular retardation (or*), which displays microphthalmia and optic nerve aplasia (Burmeister et al., 1996; Truslove, 1962). The phenotypic analysis of *or* mutants, as well as the examination of human patients with hereditary microphthalmia, revealed an essential role for *Vsx2* in neuro-epithelial proliferation and bipolar cells differentiation (Bar-Yosef et al., 2004; Burmeister et al., 1996; Ferda Percin et al., 2000). Subsequent studies indicated that, during optic cup formation, *Vsx2* is a key factor in the binary decision between neural retina and retinal-pigmented epithelium (RPE) lineages. Genetic studies in mice and chick revealed that *Vsx2* acts, downstream of the neural retina inducing ligands (i.e. FGFs), as a repressor of *Mitf* and *Tfec* genes and hence of the RPE identity (Horsford et al., 2005; Nguyen & Arnheiter, 2000; Rowan et al., 2004).

A few years after *Vsx2* identification, a closely related paralog, *Vsx1*, was reported in several vertebrate species (Chen & Cepko, 2000; Chow et al., 2001; Passini et al., 1997). The proteins encoded by these paralogous genes have similar domains’ architecture, including well-conserved paired-like homeodomain and CVC (Chx10/Vsx-1 and ceh-10) regulatory modules, and share biochemical properties, binding with high affinity to the same DNA sequence motif “TAATTAGC” (Capowski et al., 2016; Dorval et al., 2005; Ferda Percin et al., 2000; Héon et al., 2002). Although both genes display partially overlapping expression patterns in the retina, *Vsx2* precedes *Vsx1* expression in undifferentiated progenitors in all vertebrate models analysed. Furthermore, once retinal precursors exit the cell cycle, they are expressed in complementary sets of differentiated bipolar cells. Thus, *Vsx1* is restricted to different types of ON and OFF cone bipolar cells in mice, and *Vsx2* to S4 bipolar and Müller cells in zebrafish (Ohtoshi et al., 2004; Shi et al., 2011; Vitorino et al., 2009). In contrast to *Vsx2, Vsx1* seems to have a minor contribution to retinal specification in mammals. A single case of sporadic microphthalmia has been associated to *Vsx1* mutation in humans (Matías-Pérez et al., 2018), and its mutation in mice does not affect early retinal development even in a *Vsx2* mutant background (Chow et al., 2004; Clark et al., 2008). However, *Vsx1* mutation has been linked to inherited corneal dystrophies in humans, and is associated to abnormal electroretinogram (ERGs) recordings either in mice or in patients (Chow et al., 2004; Héon et al., 2002; Mintz-Hittner et al., 2004).

Despite all these advances on the developmental role of *Vsx* genes, many questions remain open. A fundamental issue is to understand to which extent *Vsx* gene functions are conserved across vertebrates. Previous morpholino studies in zebrafish have shown that *vsx2* knockdown results in microphthalmia and optic cup folding defects (Barabino et al., 1997; Gago-Rodrigues et al., 2015; Vitorino et al., 2009). However, these findings have not been validated using knockout lines, neither the role of *vsx1* and *vsx2* in fate determination and bipolar cells differentiation has been sufficiently explored in teleost fish.

To gain insight into the universality and diversity of *Vsx* functions, we have generated zebrafish mutants for *vsx1* and *vsx2* harbouring deletions within the homeodomain-encoding exons. Surprisingly, eye morphology and size appear normal either in the individual or in the double *vsx1/vsx2* mutants, thus indicating that *vsx* genes are not essential to initiate retinal development in zebrafish. The absence of early retinal malformations facilitates the phenotypic analysis of the mutants at later embryonic and larval stages. Defects in the visual background adaptation (VBA) reflex are observed in *vsx1* mutant, and appear enhanced in double mutant larvae, suggesting partial or complete blindness. Analysis of ERG responses confirms vision loss, showing that the amplitude of the b-wave recordings is reduced in *vsx1* mutants, and absent in double mutants. Interestingly, a single wild type copy of *vsx1* is sufficient to prevent VBA and ERG defects, indicating that *vsx2* loss of function can be compensated by *vsx1*. The analysis of neuronal-specific markers confirmed that retinal progenitors fail to differentiate into bipolar cells in double mutant embryos. Instead, we show that precursors at the inner nuclear layer (INL) can remain proliferative, undergo apoptosis, or be rerouted towards other retinal lineages, particularly differentiating as Müller glial cells. Finally, we investigate whether transcriptional adaptation (El-Brolosy et al., 2019) may compensate for *vsx1/vsx2* loss-off-function during retinal specification. The transcriptomic analysis of core components of the retinal specification GRN do not support a transcriptional adaptation mechanism in *vsx1/vsx2* mutants, rather suggesting that the network robustness is by itself sufficient to sustain early eye development even in the absence of *vsx1* and *vsx2* function. In summary, whereas our work shows a conserved role for *Vsx* genes during bipolar cell differentiation, also indicates that their hierarchic weight within the eye GRNs varies considerably across vertebrate species.

## RESULTS

### Zebrafish Vsx double mutants show normal eye size but affected lamination of the retina

Despite the additional round of genome duplication occurring in the teleost lineage after the split with sarcopterygians (Meyer & Schartl, 1999), a single copy of both *Vsx1* and *Vsx2* was retained in zebrafish. In order to investigate the role of Vsx transcription factors during visual system formation in zebrafish, we generated mutants for both paralogs using CRISPR/Cas9. To optimize the generation of null animals, we targeted conserved regions encoding for the DNA binding domain of the proteins in their corresponding loci at chromosome 17 (Fig 1a). We generated a 245bp deletion in *vsx1* encompassing exon3, intron3 and exon4 of the gene (*vsx1*Δ245). This mutation results in an in-frame deletion of 53 amino acids by the removal of 159pb from exon3 (54bp) and exon4 (105bp). In the case of *vsx2*, a 73bp deletion was generated in exon 3 (*vsx2*Δ73). This mutation deletes 24 amino acids of the core DBD of the protein and generates a premature stop codon in that domain. Both deletions can be easily screened by PCR with primers flanking the mutation sites. At 2-week post fertilization, no obvious macroscopic defects were observed in the visual system of either homozygous single mutants (i.e. *vsx1*Δ245 or *vsx2*Δ73) or homozygous double mutants *vsx1*Δ245; *vsx2*Δ73 (here termed *vsxKO*), which appeared normal in shape and size (Fig S1, FigS2a-d). Homozygous single mutants, and even animals harbouring a single wild type copy either of *vsx1* (*vsx1*Δ245+/-, *vsx2*Δ73-/-) or *vsx2* (*vsx1*Δ245-/-;*vsx2*Δ73+/-) reached adulthood and were fertile. However, double mutant larvae (*vsx1*Δ245 -/-; *vsx2*Δ73 -/-) died at around 3-week post fertilization, with the exception of a single unfertile escaper reaching adulthood (1 out of more than 152 larvae raised). For further analyses, double mutant embryos and larvae were obtained each generation by in-crossing of *vsx1*Δ245+/-;*vsx2*Δ73-/- or *vsx1*Δ245-/-;*vsx2*Δ73+/- animals. Once the proper recombinants were obtained, heterozygous lines maintenance was facilitated by the linkage between *vsx1* and *vsx2* mutant alleles, which tend to segregate together due to their proximity (10,6 Mb) in chromosome 17.

**Figure 1.**
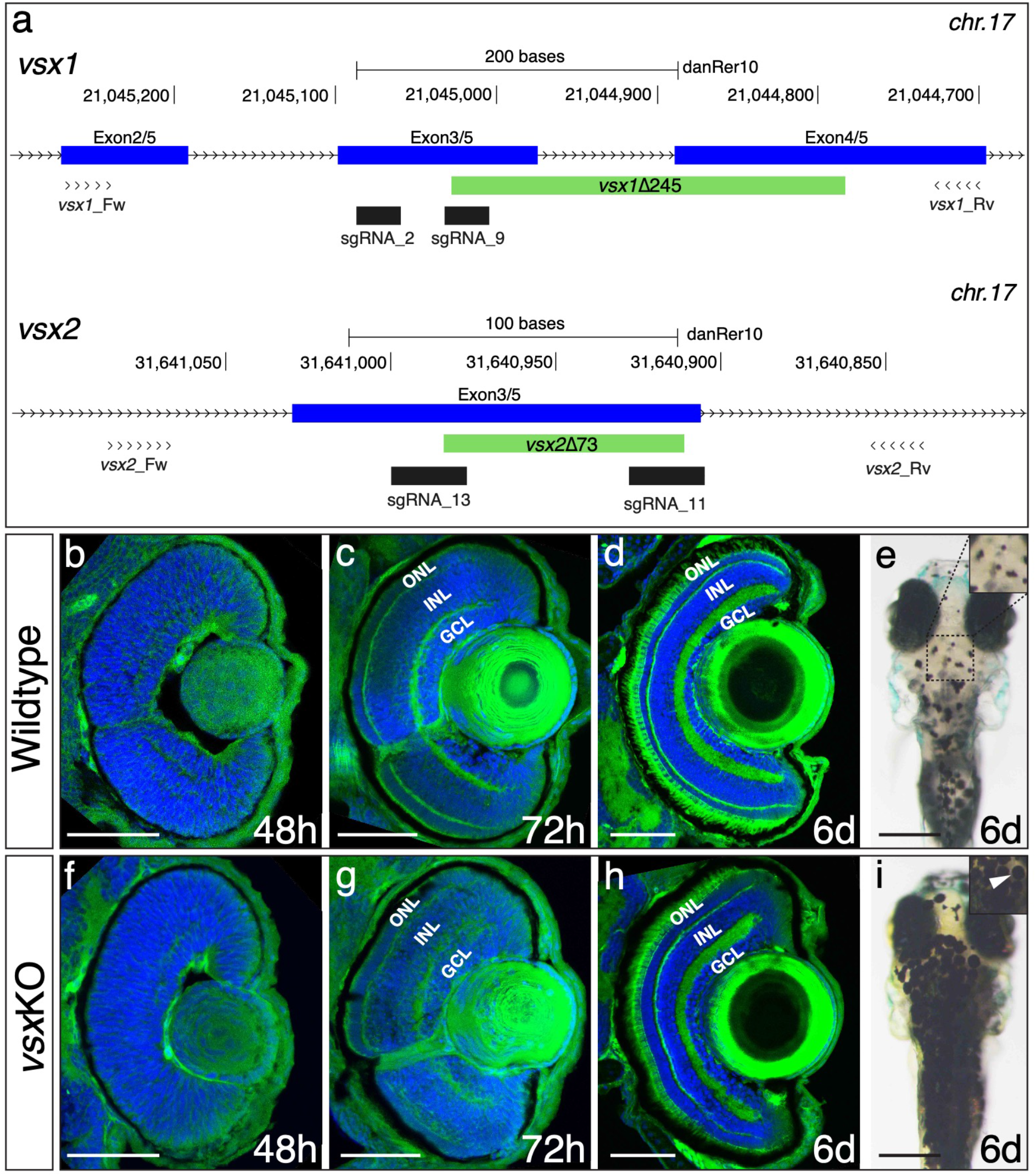
DNA binding domain deletion of vsx genes affect neural retina formation and disrupt VBA reflex. a. CRISPR/Cas9 DNA editing tool was used to generate deletions (green box) in the highly conserved DBD from *vxs1* (top) and *vsx2* (bottom) TFs. Blue boxes represent gene exons, black boxes the location of sgRNAs used to guide Cas9 endonuclease and primers for screening are depicted as opposing arrowheads. b-d and f-h. Histological sections stained with nuclear marker DAPI and phalloidin-Alexa488 for actin filaments from WT (b-d, n≥8) and *vsx*KO central retinas (f-h, n≥10) at 48hpf (b, f), 72hpf (c, g) and 6dpf (d, h). e, i. Head dorsal view from 6dpf WT (e) and *vsx*KO (i) larvae with insets showing their pigmentation pattern (white arrowhead). ONL: outer nuclear layer, INL: inner nuclear layer, GCL: ganglion cell layer, hpf: hours post-fertilization, dpf: days post-fertilization. Scale bar in b-d and f-h: 50μm, scale bar in e and i: 500μm.

Histological sectioning of mutant retinas at 48hpf showed a small delay in the formation of the inner plexiform layer (IPL), but no obvious macroscopic optic cup malformations when compared to WT (Fig 1b, f). At 72hpf, both the outer plexiform layer (OPL) and the IPL appeared less organized in the double mutant retinas, which showed discontinuities/fenestrae (Fig1c, g). At 6dpf, double mutant larvae showed all the layers of a normal retina, but the thickness of the outer (ONL) and inner (INL) nuclear layers was significantly increased and reduced respectively, when compared to siblings (Fig 1d, h; Fig S2e, h, i). In addition to retinal layer formation defects, *vsxKO* fish presented expanded pigmentation in skin melanocytes even when exposed to bright light for 20 minutes (Fig S2a-d). This phenomenon is indicative of an impaired visual background adaptation (VBA) reflex, and is often associated with blindness in zebrafish (Fleisch & Neuhauss, 2006).

### Visual function is impaired in *vsx1*Δ245-/- and vsx1Δ245-/-, vsx2Δ73-/- double mutants

To test the visual performance of the vsx1Δ245-/-, vsx2Δ73-/- mutants; ERG recordings were obtained from WT and *vsx* mutants at 5 dpf (Fig 2). Zebrafish retina becomes fully functional at 5 dpf with the exception of late maturing rods (Bilotta et al., 2001) and thus, the recorded field potentials were mainly contributed by cones. Wild type larvae show a standard ERG response to light flash, characterized by a large positive b-wave representing the depolarization of ON bipolar cells (Fig 2a), which also masks the initial a-wave generated by photoreceptor (PR) hyperpolarization. Representative recordings from larvae harboring different Vsx genotypes are shown in Figure 2a. We found that vsx2Δ73-/- ERG response (green curve) was similar to the WT recording (blue curve). However, recordings in *vsx1*Δ245-/- larvae showed a reduced b-wave compared to WT or vsx2Δ73-/- larvae. From the 36 double mutant larvae recorded in total, 10 of them still showed a b-wave, though reduced in comparison to *vsx1*Δ245-/- mutants, and much smaller than WT recordings. Moreover, in the remaining 26 double mutants recorded only the negative a-wave but no the b-wave (gray curve) was detected, suggesting that bipolar cells differentiation and/or function might be compromised. Statistical analysis of the average amplitude showed that the b-wave is significantly decreased in both *vsx1*Δ245-/- single and vsx1Δ245-/-, vsx2Δ73-/- double mutants in comparison to WT at all tested light intensities (Fig 2b). In addition, the b-wave response amplitude in the double mutant was significantly reduced compared to *vsx1*Δ245-/- single mutants (Fig 2b). These measurements are in line with our previous observation indicating that double mutant retinas are more affected at the cellular level than single *vsx1*Δ245-/- animals (Fig S2).

**Figure 2.**
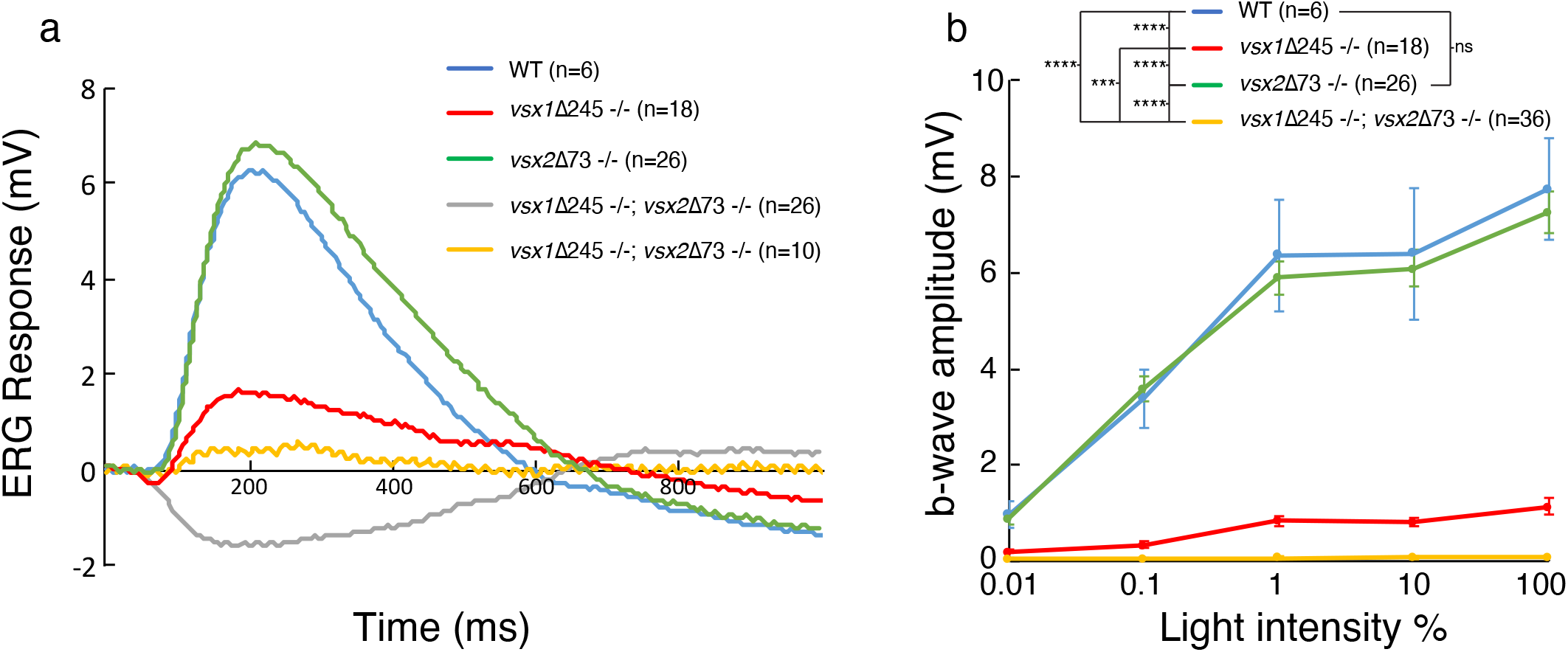
ERG response is reduced in *vsx*KO larvae. a. Representative ERG tracks at maximum light intensity from WT (blue), *vsx1*Δ245-/- (red), *vsx2*Δ73-/- (green) and *vsx1*Δ245-/-; *vsx2*Δ73-/- double mutants (grey and yellow) at 5dpf. For *vsx*KO larvae, two typical recordings are shown (grey and yellow tracks). b. Averaged ERG b-wave amplitudes from WT (blue), *vsx1*Δ245-/- (red), *vsx2*Δ73-/- (green) and *vsx1*Δ245-/-; *vsx2*Δ73-/- (yellow) larvae. No significant differences were observed between WT and *vsx2*Δ73-/- samples. *vsx1*Δ245-/- and *vsx1*Δ245-/-; *vsx2*Δ73-/- mutants produce a significant reduction of the ERG b-wave amplitude compared with both WT and *vsx2*Δ73-/- larvae throughout all light intensities tested (*p<0.01). Data are shown as mean±SEM. In a and b, *vsx1*Δ245-/- (red tracks) represents both *vsx1*Δ245-/- and *vsx1*Δ245-/-; *vsx2*Δ73+/- genotypes, while *vsx2*Δ73-/- (green tracks) represents both *vsx2*Δ73-/- and *vsx1*Δ245+/-; *vsx2*Δ73-/- genotypes. Data were collected from 5 independent experiments. For statistical comparison, one way ANOVA test was used. ms: milliseconds, mV: millivolts.

To quantitatively characterize eye performance, optokinetic response (OKR) recordings (Rinner et al., 2005) were obtained for WT and *vsx* mutant fish (Fig S3). To investigate the role of Vsx transcription factors at the behavioral level, eye movement velocity was recorded at 5dpf in WT and Vsx mutant fish. We measured eye velocity varying different parameters of the moving stimuli, such as contrast (contrast sensitivity, Fig S3a), frequency (spatial resolution, Fig S3b) and angular velocity (temporal resolution, Fig S3c). In all conditions tested, we observed a significant reduction in eye velocity for *vsx1*Δ245-/- single and vsx1Δ245-/-, vsx2Δ73- /- double mutants when compared with vsx2Δ73-/- larvae and WT controls (repeated measurement, ANOVA p<0.001). Taken together these physiological recordings confirmed significant sight impairment in *vsx1* mutants, a phenotype that is further aggravated by *vsx2* loss in *vsxKO* double mutants.

### Extended proliferation wave and INL cell death in vsx1Δ245-/-, vsx2Δ73-/- double mutant retinas

As *vsx1*Δ245-/-, *vsx2*Δ73-/- double mutants (i.e. *vsxKO*) showed stronger retinal architecture and visual defects than other *vsx* mutant combinations, we decided to focus further phenotypic analyses on them. To assess whether our observations on the increased thickness of the ONL and the decreased width of the INL (Fig S2) correlate with a proliferation and/or cell death unbalance, we examined both parameters in *vsxKO* fish. To investigate proliferation defects, we quantified the number of phosphohistone H3 positive (PH3+) cells in the retina of wild type and *vsxKO* animals throughout the lamination process: i.e. at 48, 60, and 72hpf (Fig 3a-f, m). At 48hpf, no difference in the number of PH3+ cells was observed between WT and *vsxKO* retinas (Fig 3a, d, m). However, at 60hpf, when the proliferation wave has largely finished in WT eyes, double mutant retinas continued to divide and showed a significant increase in PH3+ cells, particularly in the outer and peripheral regions (Fig 3b, e, m). Later on, at 72hpf, PH3+ cells were only detected in the CMZ and no significant difference in the number of proliferative cells was detected between WT and *vsxKO* retinas (Fig 3c, f, m). To test if cell death may account for the reduced INL width observed in double mutants (Fig S2h, i), we stained retinal cryosections with anti-cleaved caspase3 (C3) antibodies to detect cells that undergo apoptosis at different time points (Fig 3g-l, n). At 60hpf, C3 positive cells (C3+) could be observed rarely in WT or *vsxKO* retinas (Fig 3g, j, n). However, at both 72 and 96hpf, a significant increase in the number of apoptotic C3+ cells was detected in double mutants compared to WT (Fig 3h, i, k, l). Apoptotic cells concentrated mainly in the INL layer of the retina (Fig 3k, l, n), suggesting that cell death within this layer may contribute to the decreased thickness observed in *vsxKO* retinas. We also observed a few apoptotic C3+ cells in the ganglion cell layer (GCL) in *vsxKO* embryos (Fig 3k, l) suggesting than the survival of these cells may be compromised. To investigate this point, we decided to analyze the integrity of the retinal ganglion cells’ (RGCs) projections to the optic tectum by injecting DiI and DiO tracers in WT and *vsx* double mutant eyes at 6dpf (Movie S1). No obvious differences in retino-tectal projections were detected between WT and double mutant larvae, indicating that the RGCs are not affected in *vsxKO* retinas compared to control animals.

**Figure 3.**
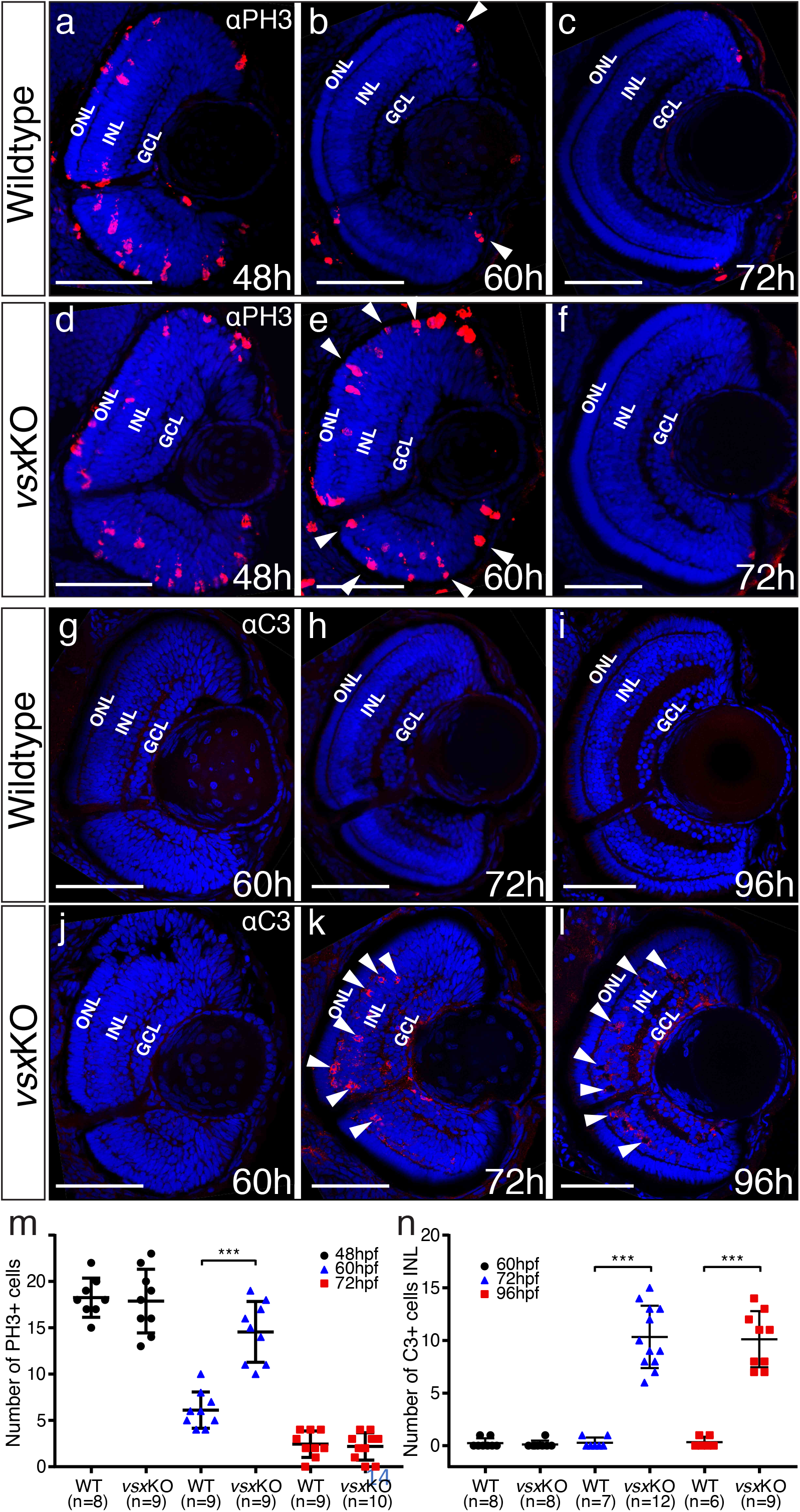
Mitosis and apoptosis markers expression are increased in *vsx*KO retinas. a-f. Phospho-histone H3 (PH3) antibody staining reveals cell divisions in central retina cryosections from WT (a-c) and *vsx*KO (d-f) samples at 3 different developmental stages (48, 60 and 72hpf). Increased PH3 staining was observed in *vsx*KO retinas at 60hpf (white arrowheads in e) compared to WT samples (white arrowheads in b). g-l. Caspase-3 (C3) antibody staining was used to evaluate cell death in central retina cryosections from WT (g-i) and *vsx*KO (j-l) samples at 3 different developmental stages (60, 72 and 96hpf). Aberrant C3 staining was observed in *vsx*KO retinas at 72 and 96hpf (white arrowheads in k and l) compared to WT samples (h and i). m. Quantification of PH3 positive cells in WT and *vsx*KO retinas at different stages. Significant increase in PH3 positive cells was observed in *vsx*KO samples at 60hpf compared to WT (***p<0.0001) but no significant changes were observed at other stages analysed (48 and 72hpf). n. Quantification of C3 positive cells in WT and *vsx*KO retinas at different stages. Significant increase in C3 positive cells was observed in *vsx*KO samples at 72 and 96hpf compared to WT (***p<0.0001), but no change was observed at 60hpf. ONL: outer nuclear layer, INL: inner nuclear layer, GCL: ganglion cell layer, hpf: hours post-fertilization. Scale bar in a-l: 50μm.

### Abnormal cell fate specification in the retina in *vsxKO*

Our results indicated that, in contrast to *Vsx*2 early requirement in the mouse (Burmeister et al., 1996), *vsx* genes are not essential for the early specification of the neural retina in zebrafish (Fig 1, Fig S1). This fact facilitated the analysis of cell fate choices in *vsxKO* embryos. Although all retinal layers are present in double mutant animals (Fig S2), the identity of the cells within these layers required further investigation. To examine cell fate acquisition in the INL and ONL of mutant retinas, fluorescent antisense probes for specific markers of PRs (*prdm1a*), bipolar (*pkcb1*), amacrine (*ptf1a*) and Müller glia cells (*gfap*) were examined at 72hpf (Fig 4).

**Figure 4.**
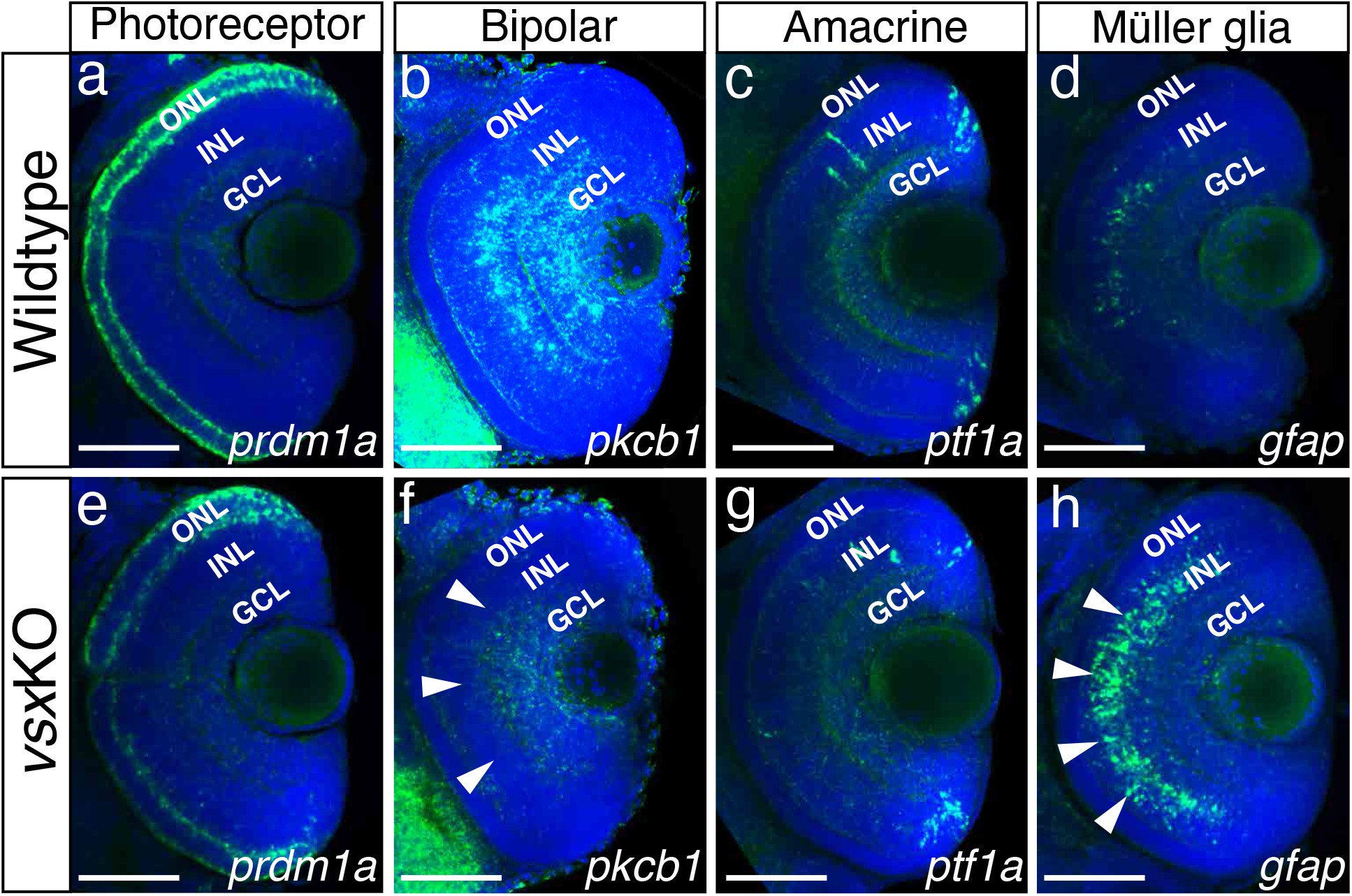
Altered expression of Bipolar and Müller glia cell markers in 3dpf *vsx* mutant fish. a-h. Confocal sections from *in toto in situ* hybridization experiments using specific fluorescent probes to label different cell types in wildtype and *vsx*KO retinas at 72hpf. No clear differences in the expression of the photoreceptor marker *prdm1a* were observed in ONL of wildtype (a) and mutant samples (e). Bipolar cell marker *pkcb1* expression (b, f) is considerably reduced in the INL of *vsx*KO mutant retinas (f, white arrowheads) compared to wildtype (b). Similar expression of the amacrine cell marker *ptf1a* is observed in the INL from wildtype (c) and *vsx*KO (g) retinas. Increased expression of the Müller glia cell marker *gfap* (d, h) is observed in the INL of double mutant samples (h, white arrowheads) compared to wildtype (d) retinas. ONL: outer nuclear layer, INL: inner nuclear layer, GCL: ganglion cell layer. Scale bar in a-h: 50μm.

#### ONL/photoreceptors specification

Prdm1a (or Blimp1) is a transcription factor that has been shown to play an early role in the specification of PR identity, mainly by the suppression of bipolar cell fate genes, including *vsx2* (Brzezinski et al., 2010; Katoh et al., 2010). Conversely, *vsx2* acute knockdown by electroporation in the postnatal mouse retina triggers a bipolar to rod fate shift (Goodson et al., 2020; Livne-Bar et al., 2006). In this study, the comparative analysis of the transient marker *prdm1a* (Wilm & Solnica-Krezel, 2005) between wild type and *vsxKO* embryos revealed similar levels of expression at 72 hpf (n=6), indicating that PRs are correctly specified in the mutants (Fig 4a, e). However, when we examined terminal differentiation markers for cones (Ab zpr-1) and rods (Ab zpr-3) at 72 and 96hpf, a delayed differentiation of both cell types was observed in double mutant embryos (Fig S4). Whereas zpr-1 and zpr-3 staining could be detected in the entire ONL in wild type fish from 72hpf on (Fig S4 a-d), in 72 hpf *vsxKO* embryos the staining was restricted to a few cells in the ventral retina and was only extended to the entire ONL at 96 hpf (n=10) (Fig S4 e-h). This result suggests that PRs’ differentiation program is delayed in the absence of *vsx* function. A prolonged period of precursors’ proliferation and/or competence could account for an increased number of PRs at larval stages, and thus for an expanded thickness of the ONL layer, as observed in double mutants at 6 dpf (Fig 1, FigS2).

#### INL/bipolar cells specification

In the zebrafish retina, *vsx1* and *vsx2* expression has been reported in complementary subsets of bipolar cells, with *vsx1* having a broader distribution and *vsx2* being restricted to a few bipolar subtypes (Vitorino et al., 2009). To analyse bipolar cell integrity in *vsxKO* embryos, we performed fluorescent *in situ* hybridizations for the bipolar cell marker protein kinase Cb1 (*prkcb1*). We found that *prkcb1* expression is very reduced, if not absent, in the INL of mutant retinas compared to WT at 72hpf (n=5) (Fig4 b, f). This result, is in agreement with our previous histological (i.e. reduced INL thickness) and electrophysiological (i.e. reduced b-wave) observations in *vsxKO* larvae, and confirms that *vsx* genes are essential for bipolar cells specification in zebrafish.

#### *INL/amacrine cells* (AC) *specification*

A detailed histological analysis of the INL architecture in wild type and *vsxKO* embryos suggested that ACs specification was not severely affected in the double mutant (Fig 1 d, h). To confirm this point, we followed the expression of *ptf1a*, a transcription factor encoding gene that is expressed in horizontal and AC and has been shown to play an essential role in their specification in the mouse retina (Fujitani et al., 2006). Using a fluorescent probe against *ptf1a*, which is expressed transiently in all types of amacrine cells in the embryonic zebrafish retina (Jusuf & Harris, 2009), we could determine that the ACs differentiation wave progresses through the central retina in wild type and *vsxKO* embryos at 48 hpf (n=5) (Figure S5). Later in development, at 72 hpf, *pft1a* expression was no longer detected in the central retina and appeared restricted to the most peripheral region, being expressed at similar levels in wild type and *vsxKO* retinas (n=10) (Fig4 c, g). This observation suggests that *vsx* genes do not play a major role for amacrine cells specification.

#### INL/Müller glia cell specification

Müller glia cell bodies are located in the INL where they provide structural and functional support to the retinal neurons (Goldman, 2014). To investigate if their differentiation occurs normally in vsx double mutants, we used a *gfap* antisense probe as glial marker (Bernardos & Raymond, 2006). We found a clear increase in the expression of *gfap* in mutant retinas compared to WT (n=5) (Fig 4d, h), suggesting that this cell type is overrepresented in *vsxKO* retinas, which may compensate for the reduction in bipolar cells observed in these animals.

In addition to their expression in the retina, Vsx transcription factors are also expressed in spinal cord interneurons (V2a and V2b cells), which are important to coordinate motor neuron activity and locomotion (Crone et al., 2008; Kimura et al., 2008). As reported here, *vsx* double mutants die around 2-weeks postfertilization. This lethality could be due by spinal cord interneuron specification defects that may restrict the movement of the animals. To examine the integrity of V2a and V2b interneurons, we label both cell types with *vsx1* and *tal1* fluorescent antisense probes, that are expressed in V2a and V2b neurons, respectively (Fig S6). No significant differences in V2a or V2b spinal cord interneurons density was observed between WT (n=6) and vsx1Δ245-/-, vsx2Δ73-/- mutant fish (n=8) at 24hpf (Fig S6e). These results suggest that V2 motoneurons are properly specified in *vsxKO* animals. In agreement with this observation, obvious swimming defects were not observed in *vsxKO* larvae.

### RNA-seq and ATAC-seq analyses of *vsxKO* reveal eye GRN robustness

The strong microphthalmia and abnormal specification of the neural retina reported in *vsx2* mutant mice (Burmeister et al., 1996; Horsford et al., 2005; Rowan et al., 2004) are in contrast to our observation that in *vsxKO* embryos/larvae neuroretinal identity is normally established and maintained. To gain insight into the molecular causes behind this discrepancy, we sought to investigate transcriptional and chromatin accessibility changes in mutant embryos during the specification of the neural retina (Fig 5). To this end, 18hpf embryo heads were collected from *vsxKO* and their wild type siblings and the rest of the tissue was used for PCR genotyping. We focused in this particular stage as it corresponds to the early bifurcation of the neural retina and RPE GRNs in zebrafish (Buono et al., 2021). To first identify changes in the cis-regulatory landscape associated to *vsx* loss of function, we examined wild type and mutant samples using ATAC-seq. This approach identified 1564 DNA regions with differential accessibility, most of them (1204) with a high fold change (i.e log2 fold change > |1.5|). They include 633 regions more accessible in the mutant with a log2 fold change > 1.5; and 571 less accessible with a log2 fold change < −1.5 (Fig 5a, b; Supplementary Dataset 1). An analysis of enriched gene ontology terms for those genes (2219) neighbouring the differentially opened regions revealed entries related to neuronal differentiation and eye development (Fig S7; Supplementary Dataset 2). This observation suggests that *vsx* genes mutation results in the deregulation of hundreds of cis-regulatory elements mainly associated to retinal genes. In contrast, at a transcriptional level the comparative analysis of mutant and wild type samples by RNA-seq revealed expression changes only in a relatively small gene set (871) (Fig 5c, Supplementary Dataset 3). This collection comprised 42 up-regulated (log2 fold change > 1.5) and 29 down-regulated (log2 Fold change < −1.5) genes, with only 3 up-regulated (*vsx1*, *znf1109* and *znf1102*) and one down-regulated transcription factor (*znf1091*) above the threshold (log2 fold change > |1.5|) (Fig 5c). This observation indicated that the identified cis-regulatory changes are only translated in subtle changes at the transcriptional level. In fact, among the 2219 genes neighbouring differentially open chromatin regions, only 5% (119) were associated to differentially expressed genes (Fig 5d). To further confirm the impact of *vsx* loss of function on the expression of core components of the neural retina GRN, we examined their levels by qPCR at 19 hpf (Fig 5e). Interestingly, in *vsxKO* embryos significant expression changes could be detected only for *rx2* and *lhx2b* (though below the threshold log2 fold change > |1.5|). In addition, the transcripts of the mutated genes *vsx1* and *vsx2* were significantly upregulated and downregulated respectively in mutant embryos at optic cup stages, as determined by qPCR at 19hpf (Fig 5e) and confirmed by ISH at 24hpf (Fig S8). Taken together, these analyses suggest that the general architecture of the retinal GRN was not significantly altered upon *vsx* genes mutation. Furthermore, they also point at *vsx* genes having a different regulatory weight for neural retina specification/maintenance in different species. To gain insight into this hypothesis, we mutated two of the three paralogs (i.e. *vsx1* and *vsx2.2*) present in the genome of the far-related teleost medaka by CRISPR-Cas9 (Fig S9). Interestingly, although the initial specification of the organ appeared normal also in medaka embryos at 4dpf, INL differentiation and eye growth was impaired at later stages 12dpf (Fig S9b-e). This is in contrast to the normal eye size observed in *vsxKO* zebrafish larvae at 19dpf (Fig S10) and confirmed the assumption of differential regulatory weight among vertebrate species for *vsx* genes.

**Figure 5.**
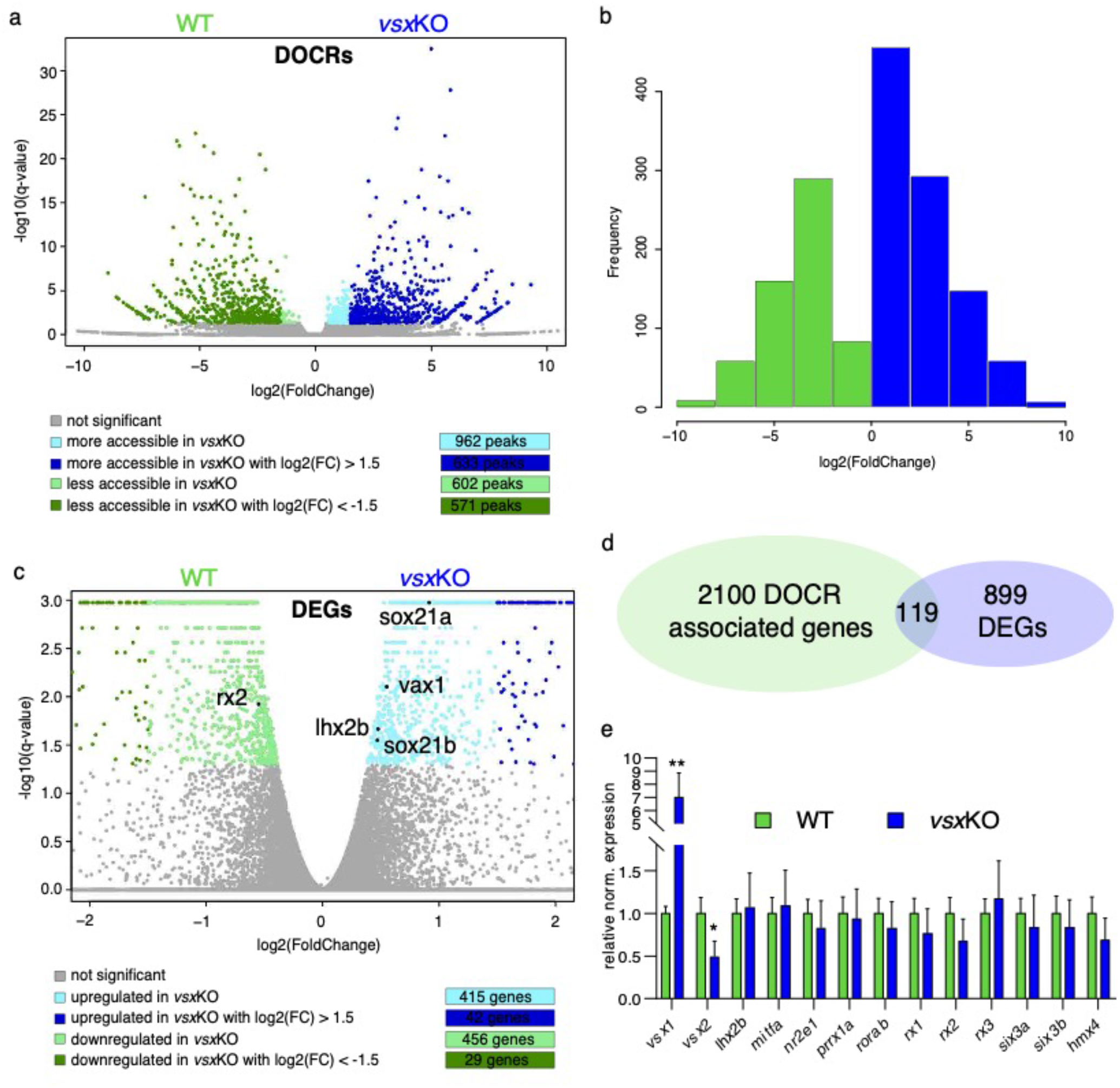
Lack of vsx TFs in the forming retina is buffered by genetic redundancy. a. Volcano plots illustrating chromatin accessibility variations upon vsx1 and vsx2 mutation in zebrafish retina at 18hpf. Each dot corresponds to an ATAC-seq peak, i.e. an open chromatin region. Grey dots indicate not significant variations, whereas coloured dots point out significant differentially open chromatin regions. b. Frequency of DOCRs’ fold change values. c. Transcriptome variations in *vsx*KO retina at 18hpf. The genes reported in the plot are the only known retinal regulators whose transcriptional levels are affected by the loss of Vsx factors, with a very modest fold change. Essentially, RNA-seq experiments did not highlight a remarkable change of the levels of the main TFs governing the retinal GRN. d. Correspondence between genes associated with DOCRs from ATAC-seq and DEGs from RNA-seq. e. qPCR of the main retinal TFs confirming the stability of the eye gene network expression after *vsx1* and *vsx2* loss (n=3). DOCR: differentially open chromatin regions, DEG: differentially expressed genes.

## DISCUSSION

In this study, we explore the universality of Vsx functions in the development of the vertebrate eye, by generating CRISPR-Cas9 mutations of the “visual system homeobox” genes *vsx1* and *vsx2* in the far related teleost models, zebrafish and medaka. Genetic analyses in the mouse, as well as the chick, had revealed two distinct functions for *Vsx* genes during eye development: an early requirement for proliferation and specification of the neural retina precursors, and a later role in the differentiation of bipolar neurons (Burmeister et al., 1996; Horsford et al., 2005; Rowan et al., 2004). These two developmental roles depend on consecutive waves of gene expression and thus can be uncoupled by genetic interference within specific developmental windows (Goodson et al., 2020; Livne-Bar et al., 2006). Moreover, in mice, Vsx biphasic activity follows a partially independent *cis*-regulatory control by enhancers active either in precursors, bipolar cells, or both (D. S. Kim et al., 2008; Norrie et al., 2019; Rowan & Cepko, 2005). Accordingly, CRISPR-mediated ablation of a distal bipolar enhancer results in the specific depletion of these cells, without leading to microphthalmia or compromising the early specification of the mouse retina (Goodson et al., 2020; Norrie et al., 2019).

Here we show that *Vsx* activity is essential for bipolar cells differentiation in teleost fish, indicating a broadly conserved role for these genes across vertebrates. This observation suggests that the genetic program controlling bipolars specification was inherited from a common vertebrate ancestor. The fact that *Vsx* homologous genes are also expressed in the visual-system of the invertebrates *Drosophila* and Cuttlefish (*Sepia officinalis*) further suggests that the function for these homeobox genes in the specification of visual interneurons may be a common theme in all metazoans (Erclik et al., 2008; Focareta et al., 2014). The absence of an earlier eye phenotype in zebrafish *vsx*KO embryos allowed us examining in detail the consequences of *vsx* loss on cell fate determination and sight physiology. Both our histological and electrophysiological analyses confirmed bipolar cells depletion in *vsx*KO retinas. We show that, unable to acquire the bipolar fate, retinal precursors follow alternative differentiation trajectories such as undergoing apoptosis, extending their proliferative phase, or differentiating as photoreceptors or Müller glia cells. A detour towards photoreceptors fate in zebrafish is in agreement with previous studies in mice showing that the Blimp1/Vsx2 antagonism controls the balance between rods and bipolar cells (Brzezinski et al., 2010; Goodson et al., 2020; Katoh et al., 2010; Wang et al., 2014). Interestingly, in *vsx*KO retinas we observed a noticeable delay in the onset of cones and rods terminal differentiation markers, *zpr-1* and *zpr-3* respectively, indicating that Vsx activity is not only required for correct fate specification, but also to determine the timing of the differentiation sequence. Arguably, more intriguing was our observation of an increased number of Müller glia cells in *vsx*KO retinas. Both glial and bipolars cells are late-born retinal types deriving from a common pool of precursors with restricted developmental potential (Bassett & Wallace, 2012; Satow et al., 2001). In mice, however, a similar increase in Müller glia cells has not been reported in experiments genetically interfering with *vsx* either postnatally (Goodson et al., 2020; Livne-Bar et al., 2006) or specifically in bipolar cells (Norrie et al., 2019). This apparent discrepancy might indicate some variations in the cell fate specification mechanisms among vertebrate species. Alternatively, the increase may have been overlooked in previous studies due to the small size of the Müller glia cell population.

Despite the severe visual impairment and retinal lamination defects we observed in *vsx*KO larvae, their eyes appear normal in shape and size and no early morphological defects are observed in the optic cup. More importantly, neuro-retinal identity seemed perfectly maintained in double mutant animals, and we did not detect any trans-differentiation of the retina into pigmented cells. This finding, which is in contrast with the microphthalmia and the neural retina specification defects observed in mice (Burmeister et al., 1996; Horsford et al., 2005; Rowan et al., 2004), may indicate that *Vsx* genes do not play an early role in the establishment and maintenance of the neural retina identity in zebrafish. Our observations are also in contrast to previous reports in zebrafish using morpholinos against *vsx2*, which show microphthalmia and optic cup malformations (Barabino et al., 1997; Gago-Rodrigues et al., 2015; Vitorino et al., 2009). A poor resemblance between morpholino-induced and mutant phenotypes has been previously described in zebrafish, with many mutations lacking observable phenotypes (Kok et al., 2015). Genetic compensation and, in particular, transcriptional adaptation (i.e. up-regulation of genes displaying sequence similarity) has been identified as the molecular mechanism accounting for genetic robustness in a number of these mutations (El-Brolosy et al., 2019; Rossi et al., 2015). However, our comparative transcriptomic analysis of *vsx*KO vs. wt. embryos does not support genetic compensation acting as a relevant mechanism at optic cup stages. We show that, despite that *vsx* loss of function results in the deregulation of hundreds of *cis*-regulatory regions associated to retinal genes, this has little impact on the expression of core components of the neural retina specification network.

Our previous analysis of transcriptome dynamics and chromatin accessibility in segregating NR/RPE populations indicated that the regulatory networks involved in the specification of the zebrafish eye are remarkably robust (Buono et al., 2021). In that study, we showed that the consensus motif 5’-TAATT-3’, which is central to the neural retina *cis*-regulatory logic, is shared by many homeodomain TFs co-expressed during retinal specification; including not only *vsx1* and *vsx2*, but also *rx1*, *rx2*, *rx3*, *lhx2b*, *lhx9*, *hmx1*, and *hmx4*. Moreover, we show evidence that these TFs may co-regulate the same genes and cooperate within the same *cis*-regulatory modules (Buono et al., 2021). According to these observations, gene redundancy appears as a more parsimonious explanation for the absence of an early phenotype in *vsx*KO embryos. This would suggest that the regulatory weight of *vsx* genes within the retina network varies across vertebrate species. Several lines of evidence support this view. (i) Other mutations in genes encoding for TFs targeting the motif 5’- TAATT-3’, such as *rx2* in medaka (Reinhardt et al., 2015) or *lhx2* in zebrafish (Seth et al., 2006) do not compromise the identity of the neural retina tissue either. (ii) Even in *vsx1/vsx2* double mutant mice, the central retina keeps the potential for differentiation into several neuronal types, indicating that other genes must cooperate in the specification of this tissue (Clark et al., 2008). In such scenario of complex epistatic interactions, it is not surprising the intrinsic variability in expressivity traditionally observed in ocular retardation mutants (Osipov & Vakhrusheva, 1983). In line with this, in *Vsx2* mutant mice has been shown that neural retinal identity defects and microphtalmia (but not bipolar cells differentiation) can be partially restored by simply deleting a cell cycle gene (Green et al., 2003). (iii) Finally, here we show that the mutation of the paralogs vsx1 and *vsx2.2* results in severe microphthalmia in medaka larvae. This finding confirms a variable role across species for Vsx genes in the specification and maintenance of the neural retina domain in vertebrates.

---

**Figure S1.**
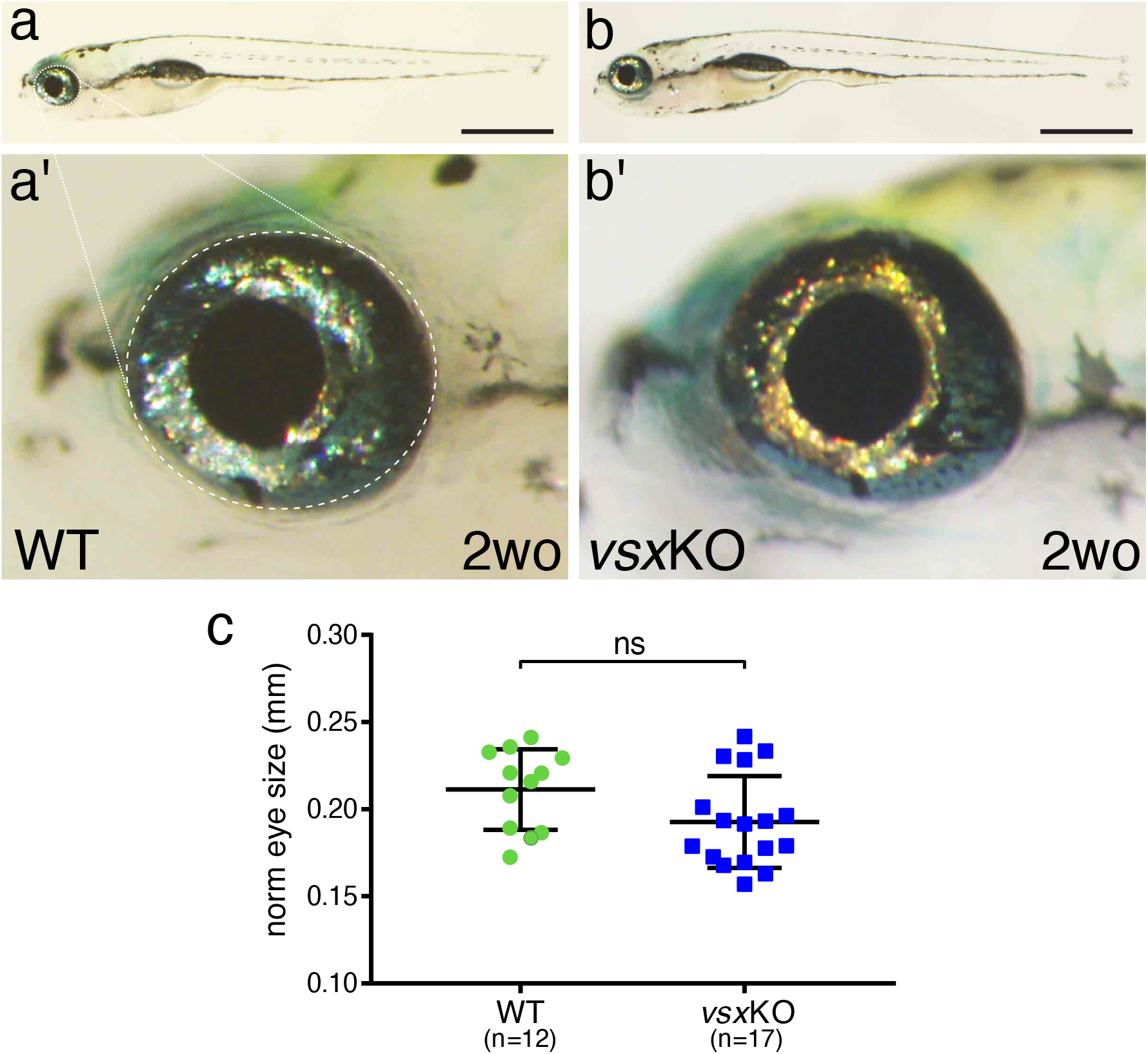
Eye size is normal in *vsx*KO juvenile fish. a, b. Lateral view of a 2-week old wildtype (a) and *vsx*KO (b) fish. Note that *vsx*KO juvenile fish show no obvious morphological differences with wildtype siblings. a’, b’. High magnification images from wildtype (a’) and *vsx*KO (b’) fish heads. No changes in eye shape or size were observed between mutant and wildtype siblings. c. Quantification of eye lateral surface (normalized by fish anteroposterior length) in 2-week old fish revealed no significant differences in eye size between wildtype (green circles) and *vsx*KO mutant fish (blue squares). Scale bar in a and b: 1mm.

**Figure S2.**
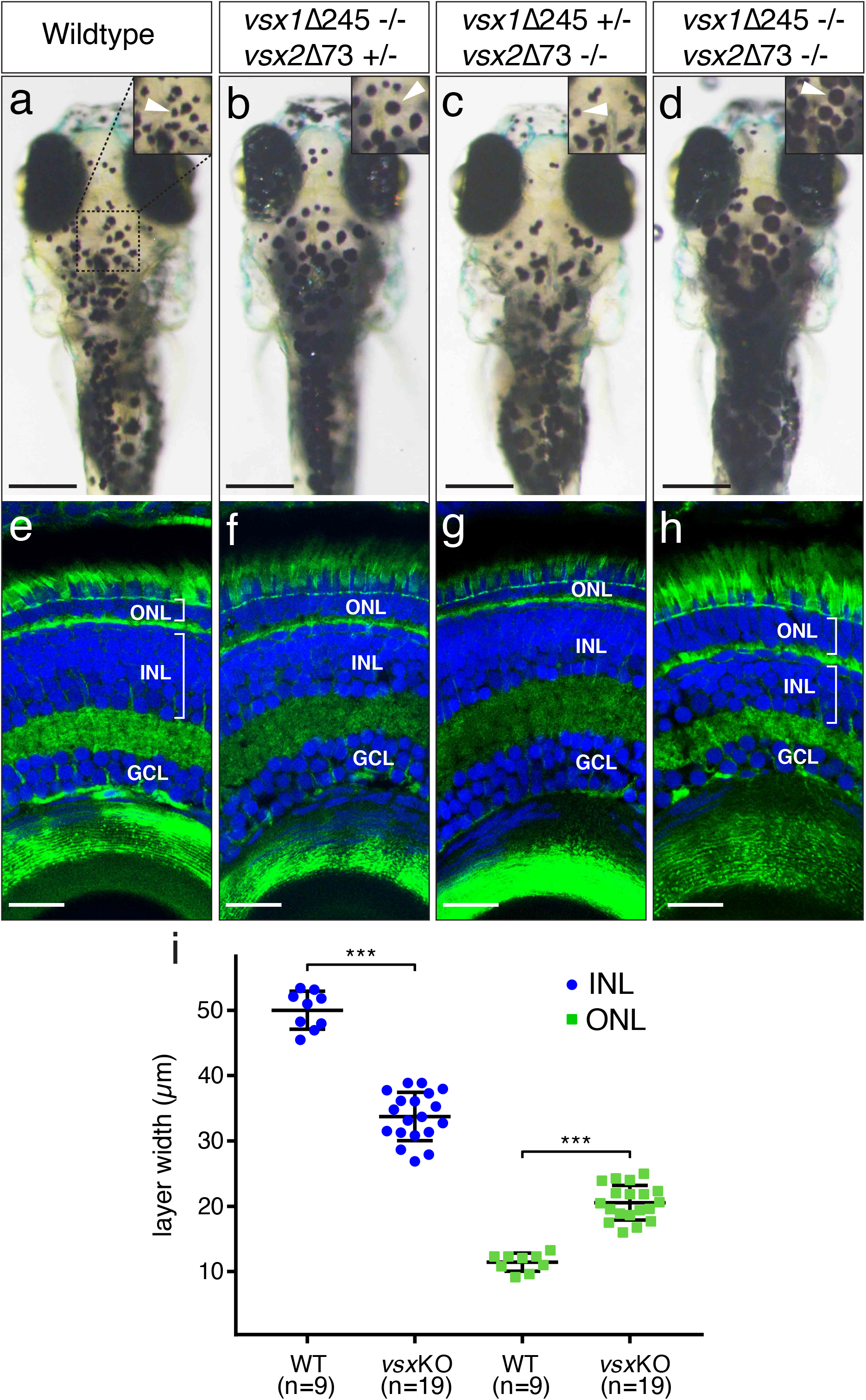
VBA and nuclear layers width are affected in *vsx* mutants. a-d. Dorsal view of wildtype (a), *vsx1*Δ245-/-; *vsx2*Δ73+/- (b), *vsx1*Δ245+/-; *vsx2*Δ73-/- (c) and *vsx1*Δ245-/-; *vsx2*Δ73-/- (d) animals at 6dpf. *vsx1*Δ245-/-; *vsx2*Δ73+/- (b) and *vsx1*Δ245-/-; *vsx2*Δ73-/- (d) fish have problems sensing background light and appear darker than wildtype (a) and *vsx1*Δ245+/-; *vsx2*Δ73-/- (c) larvae, as melanosomes are broadly distributed within melanophores (white arrowheads). e-h. Histological sections of wildtype (e), *vsx1*Δ245-/-; *vsx2*Δ73+/- (f), *vsx1*Δ245+/-; *vsx2*Δ73-/- (g) and *vsx1*Δ245-/-; *vsx2*Δ73-/- (h) eyes at 6dpf. In the central retina of *vsx*KO double mutants (h), the INL width is reduced and the ONL thickness is expanded compared to wildtype (e), *vsx1*Δ245-/-; *vsx2*Δ73+/- (f) or *vsx1*Δ245+/-; *vsx2*Δ73-/- (g) samples. DAPI and phalloidin coupled to Alexa488 were used as nuclear and cytoskeletal (actin) markers, respectively. i. Quantification of INL (blue circles) and ONL (green squares) width in central retinas at 6dpf using histological sections as in e-h panels. Significant reduction and expansion of the INL and ONL, respectively, was observed in *vsx*KO retinas when compared to wildtype (***p<0.0001). ONL: outer nuclear layer, INL: inner nuclear layer, GCL: ganglion cell layer, μm: micrometres. Scale bar in a-d: 500μm, scale bar in e-h: 20μm.

**Figure S3.**
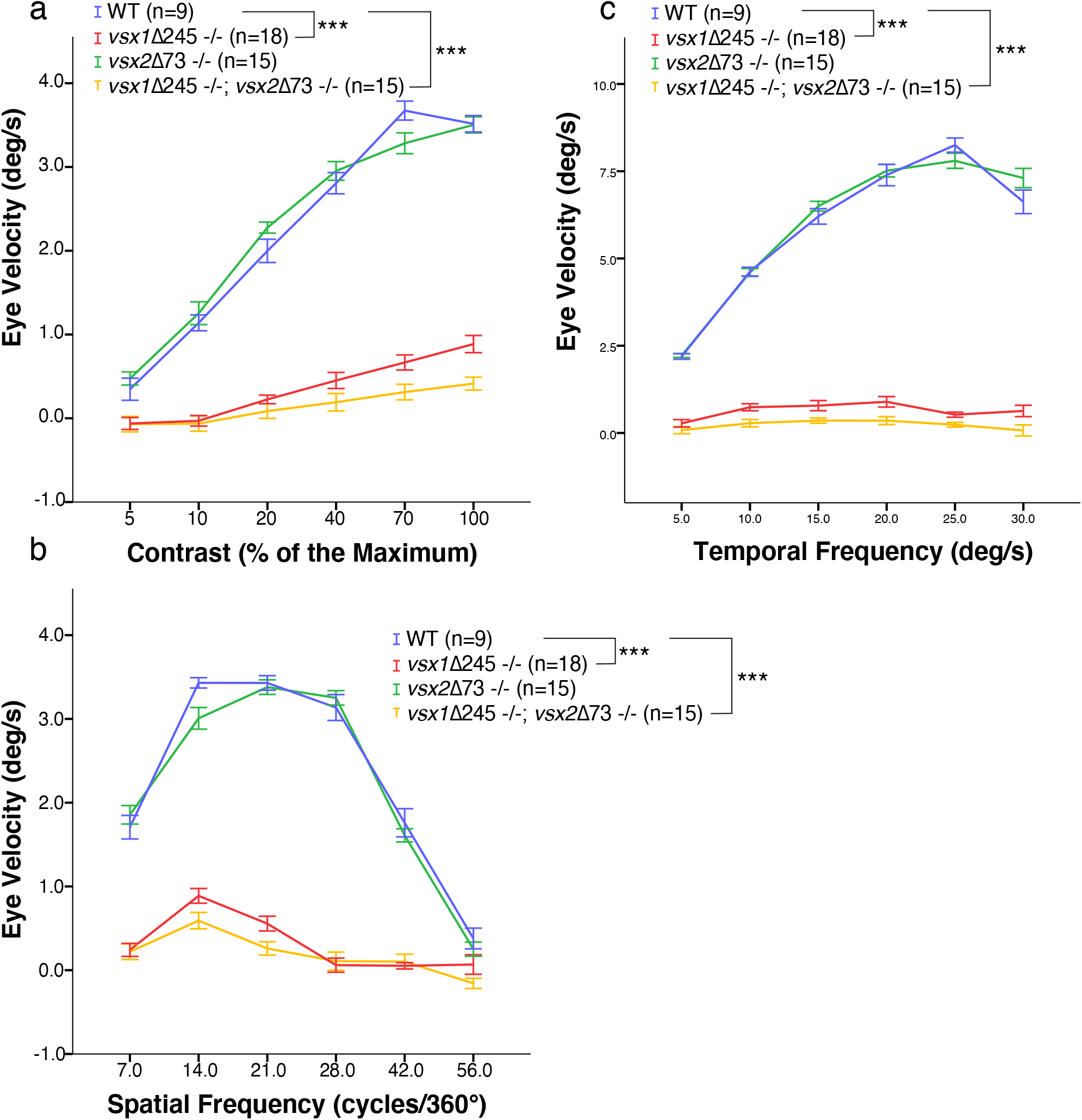
OKR measurements indicate decreased eye movement velocity in *vsx* mutants. a-c. OKR was recorded in wildtype (blue), *vsx1*Δ245-/- (red), *vsx2*Δ73-/- (green) and *vsx1*Δ245-/-; *vsx2*Δ73-/- double mutants (yellow) at 5dpf in response to different contrast (a), spatial frequency (b) and angular velocity (d). OKR is significantly reduced in *vsx1*Δ245-/- and *vsx1*Δ245-/-; *vsx2*Δ73-/- double mutant larvae in comparison to WT siblings (***p<0.001). Note that no differences in OKR were observed between WT and *vsx2*Δ73-/- single mutants or between *vsx1*Δ245-/- and *vsx1*Δ245-/-; *vsx2*Δ73-/- double mutants (p>0.05). In (a-c), *vsx1*Δ245-/- (red tracks) represents both *vsx1*Δ245-/- and *vsx1*Δ245-/-; *vsx2*Δ73+/- genotypes, while *vsx2*Δ73-/- (green tracks) represents both *vsx2*Δ73-/- and *vsx1*Δ245+/-; *vsx2*Δ73-/- genotypes. In all panels, data is shown as mean ±SEM. deg: degrees, s: seconds.

**Figure S4.**
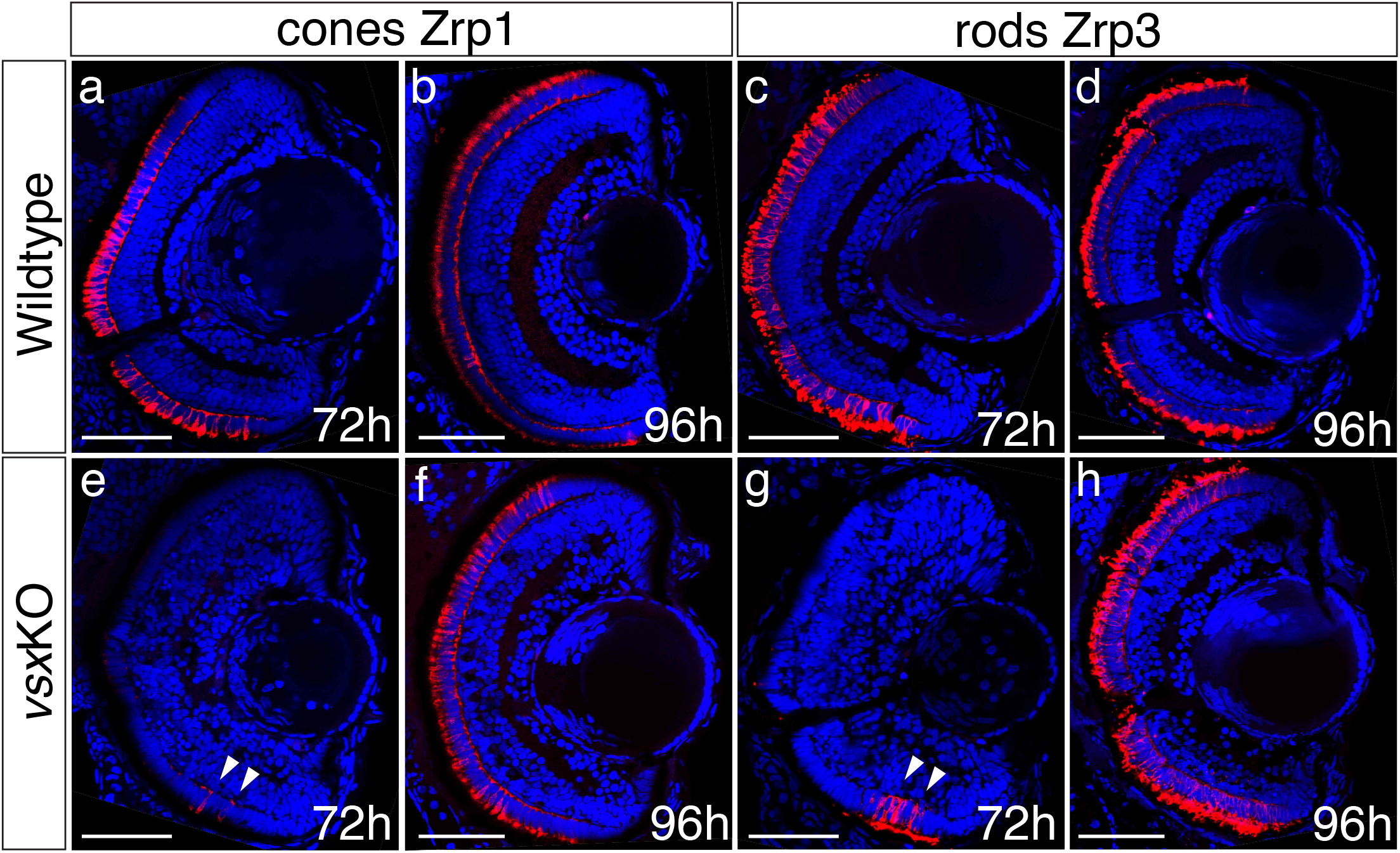
Delayed photoreceptor differentiation is observed in *vsx*KO retinas. a-h. Cryosections of wildtype (a-d) and *vsx*KO retinas (e-h) stained with photoreceptor specific antibodies and DAPI. Cones and rods were visualized by using the zpr1 (a, b, e, f) and zpr3 (c, d, g, h) antibodies, respectively. Delayed cone and rod differentiation is observed in *vsx*KO mutants (e, g) compared to wildtype (a, c) samples at 72hpf (white arrowheads in e, g). At 96hpf, both markers are expressed in similar levels in wildtype and (b, f) double mutant retinas (d, h). For all conditions tested, n≥10. Scale bar in a-h: 50μm.

**Figure S5.**
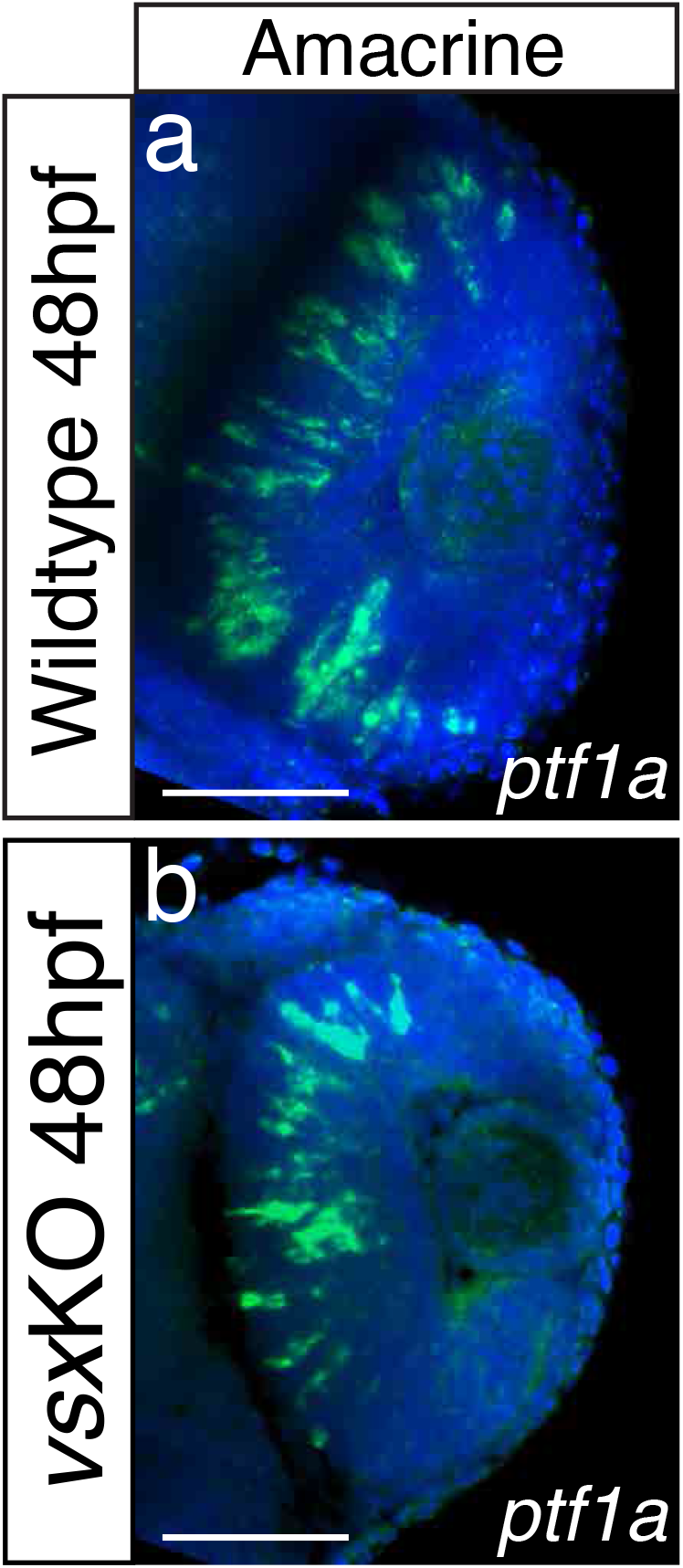
Amacrine cell marker *ptf1a* expression is not disturbed in *vsx*KO forming eyes. a-b. Confocal images from wildtype (a) and *vsx*KO (b) retinas at 48hpf after fluorescent *in situ* hybridization protocol was performed using *ptf1a* as probe to label amacrine cells. No major differences in the expression of this gene between wildtype (a) and vsxKO samples (b) were observed at this stage. Scale bar in a, b: 50μm.

**Figure S6.**
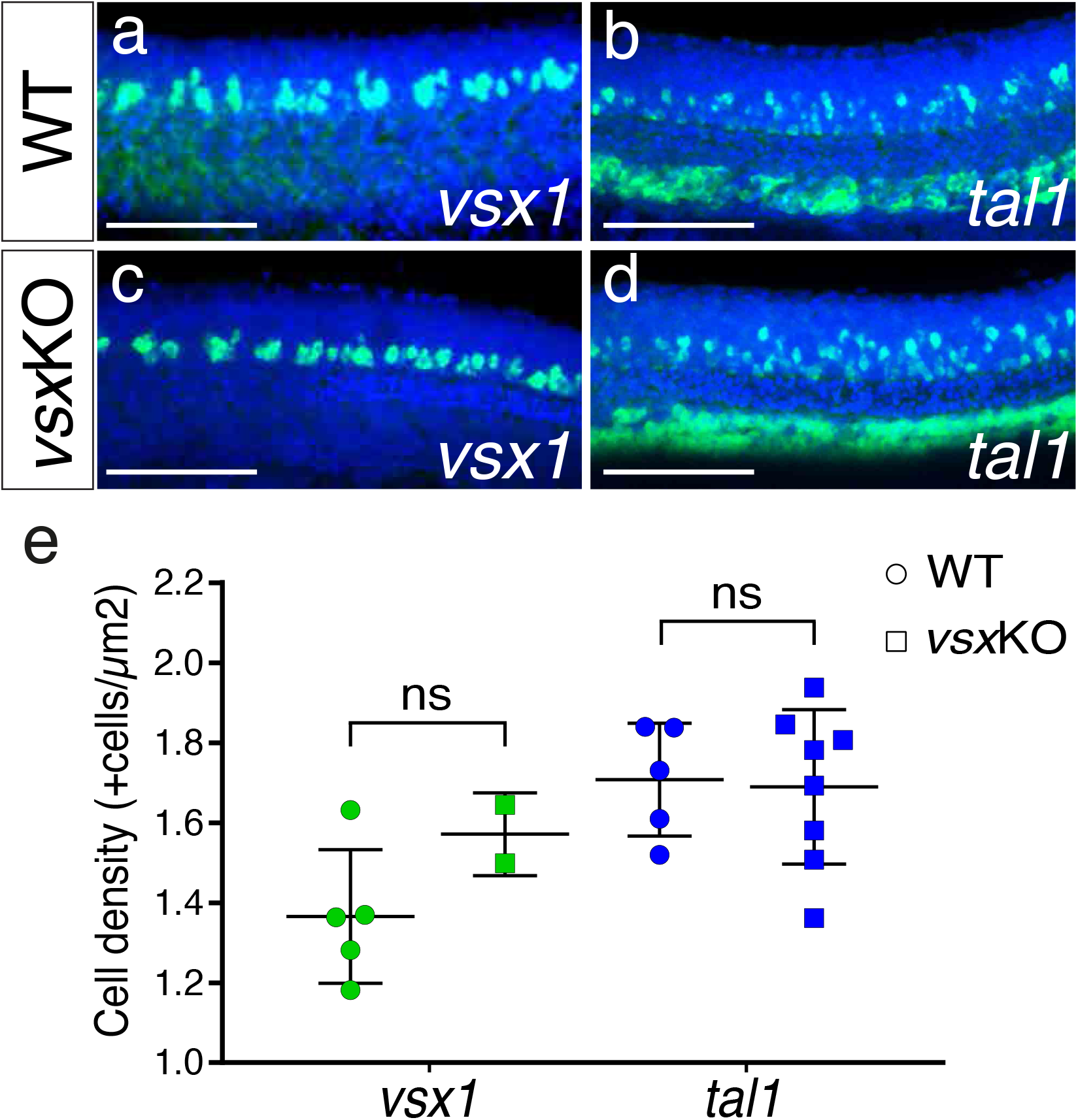
V2 spinal cord interneurons are not affected by the mutation of *vsx* TFs. a-d. Fluorescent *in situ* hybridization lateral images from 24hpf wildtype (a, b) and *vsx*KO (c, d) mutants using *vsx1* (a, c) and *tal1* (b, d) probes to visualize V2a and V2b trunk interneurons, respectively. e. The density of *vsx1* (V2a) and *tal1* (V2b) positive neurons was measured in wildtype and vsxKO mutant trunks at 24hpf. No significant changes in the number of V2a and V2b neurons was observed between wildtype and *vsx*KO mutant animals at this stage. ns: not significant. Scale bar in a-d: 100μm.

**Figure S7.**
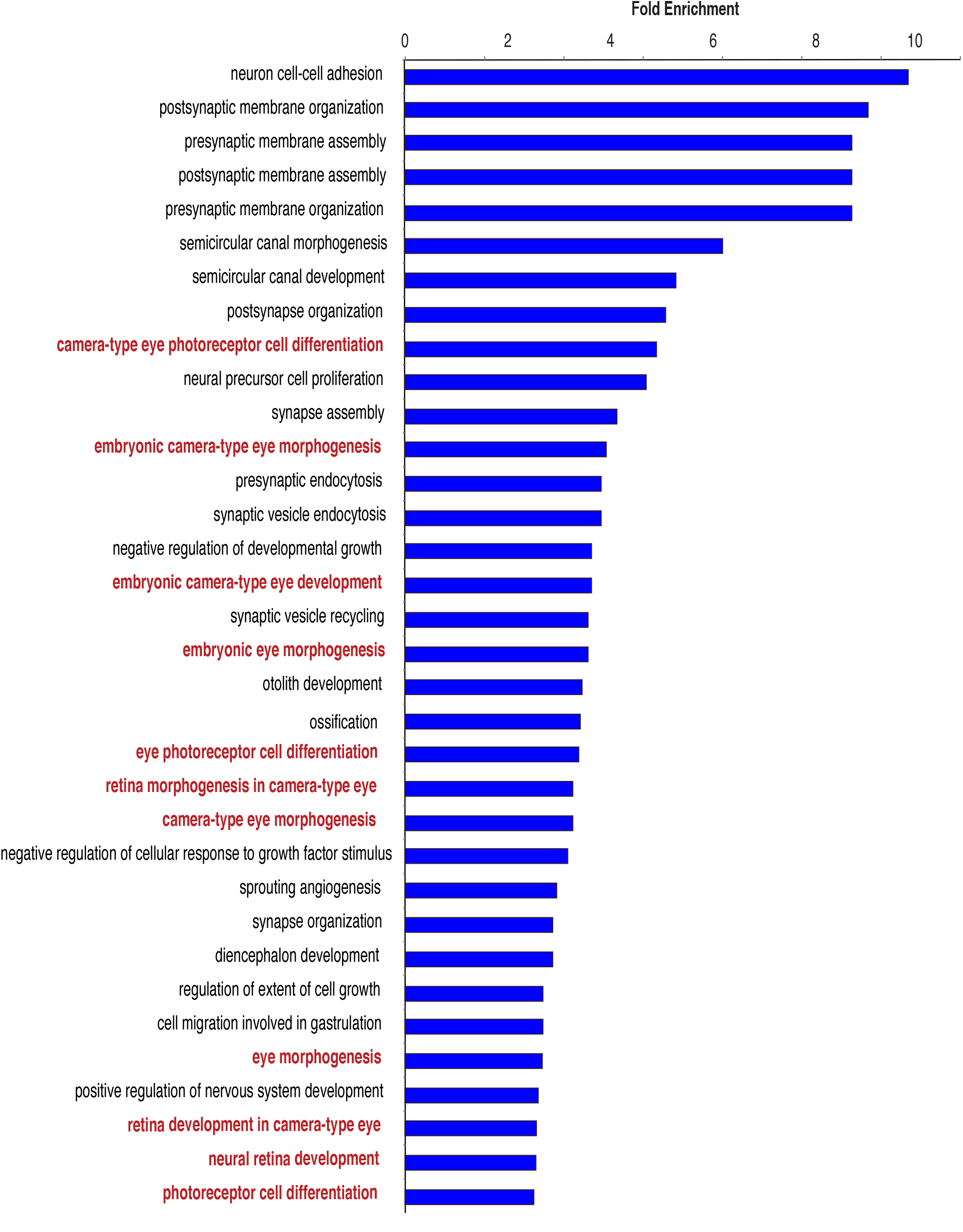
GO annotations linked to genes neighbouring DOCRs are enriched in eye development terms. “Biological process” GO terms enriched in genes associated with a more accessible DOCR in *vsx*KO embryos. GO terms directly related with eye differentiation are highlighted in red. Most GO terms not highlighted in red correspond to nervous system related processes.

**Figure S8.**
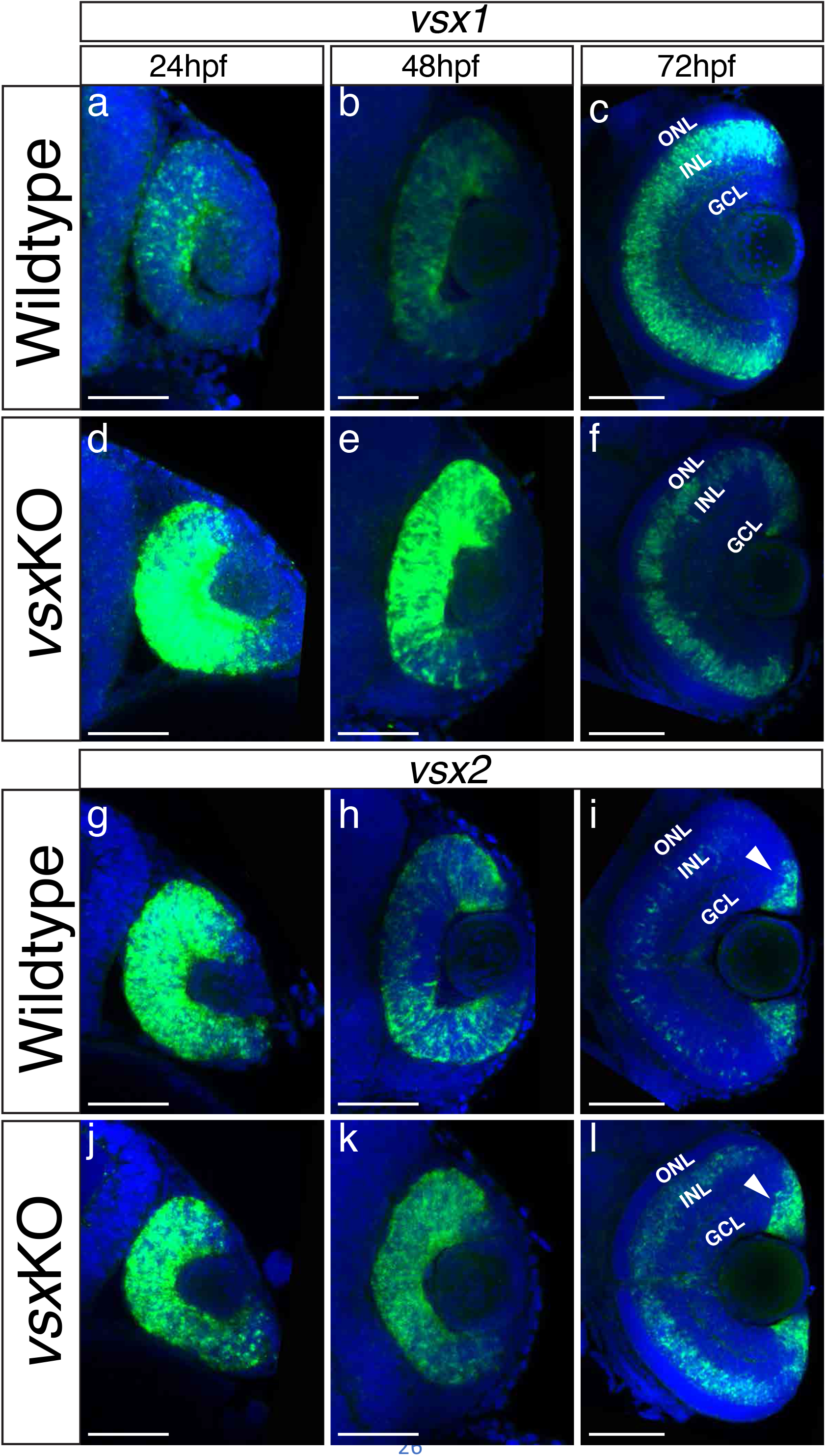
The expression pattern of *vsx* mutant transcripts is misregulated in *vsx*KO animals during retina development. a-l. Fluorescent *in situ* hybridization for *vsx1* (a-f) and *vsx2* (g-l) genes over retina formation in wildtype (a-c, g-i) and *vsx*KO (d-f) animals (j-l). In wildtype samples, *vsx2* is expressed strongly than *vsx1* in retinal precursors at earlier developmental stages (a, g) and the opposite effect is observed in INL bipolar cells at later stages (c, i). We found an overexpression of *vsx1* at 24hpf (b) and 48hpf (e) and a reduction in the INL expression at 72hpf (c, f) in *vsx* mutant retinas compared to wildtype. For *vsx2*, no major differences between wildtype and mutant retinas were detected at 24hpf (g, j), but at 48 and 72hpf we found an increment of *vsx2* in the retina and INL, respectively (h, k, i, l). Also, at 72hpf, there is a stronger expression of *vsx2* in the ciliary marginal zone (CMZ) compared to WT. hpf: hours post fertilization, ONL: outer nuclear layer, INL: inner nuclear layer, GCL: ganglion cell layer. Scale bar in a-l: 50μm.

**Figure S9.**
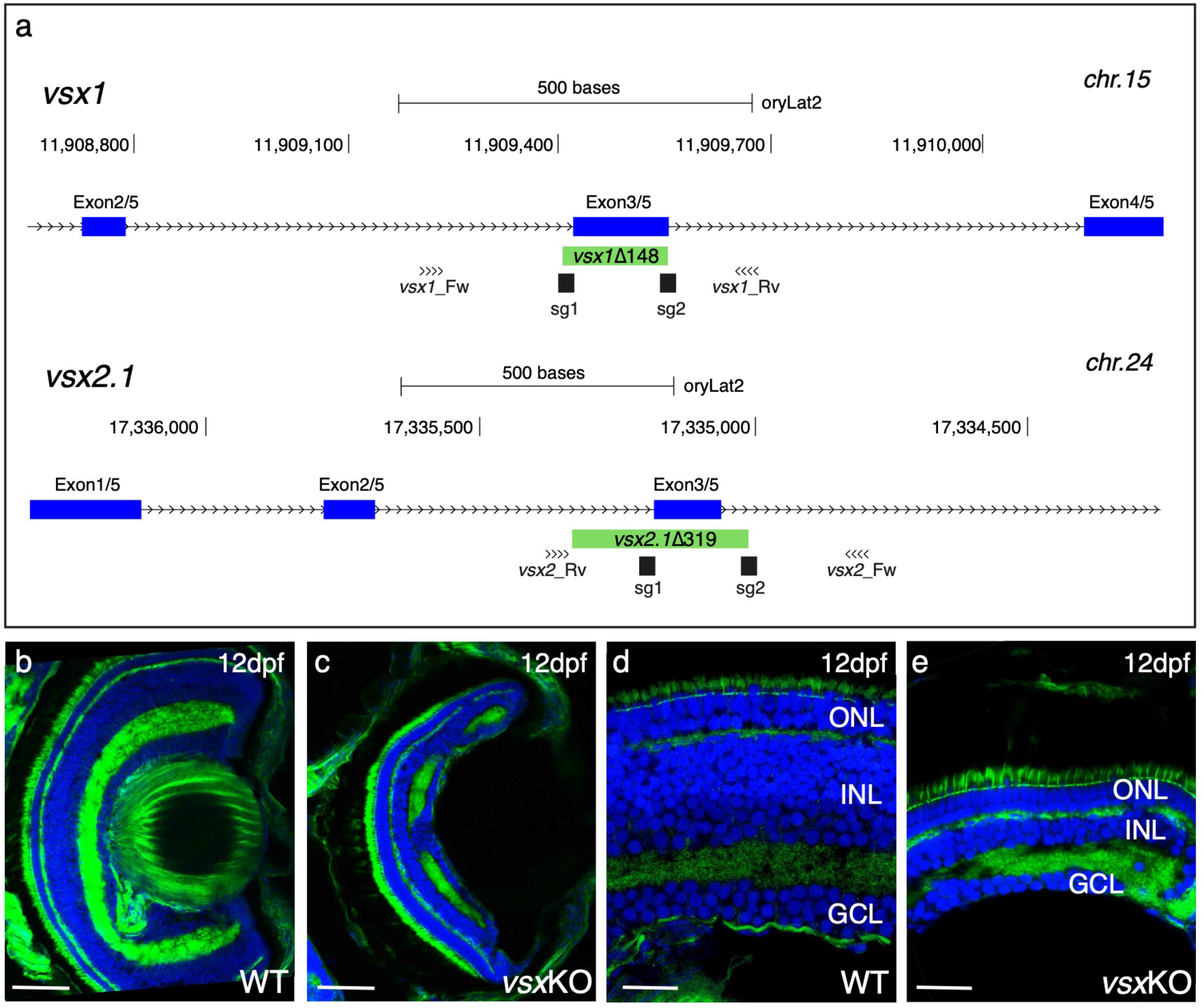
Mutation of *vsx* genes in medaka impairs INL differentiation and eye growth. a. CRISPR/Cas9 was used to eliminate (green box) the DBD from *vxs1* (top) and *vsx2.1* (bottom) TFs in medaka. Blue boxes represent exons, black boxes the location of sgRNAs used and primers for screening are depicted as opposing arrowheads. b-e. Histological sections from WT (b, d, n=4) and *vsx*KO central retinas (c, e, n=5) at 12dpf. ONL: outer nuclear layer, INL: inner nuclear layer, GCL: ganglion cell layer, hpf: hours post-fertilization, dpf: days post-fertilization. Scale bar b-c: 50μm, d-e: 20 μm.

**Figure S10.**
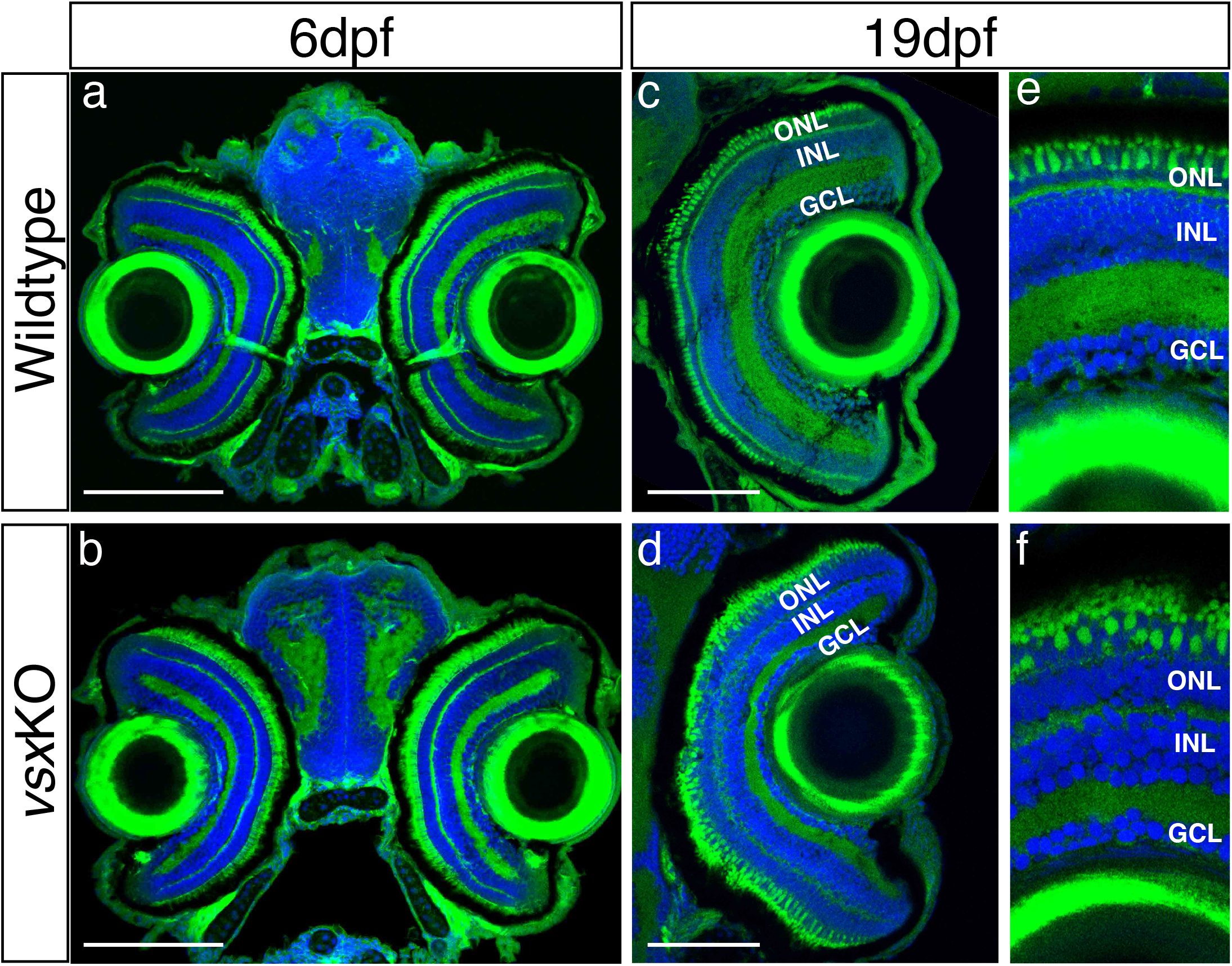
Normal eye size in *vsx*KO animals is observed at juvenile stages. a-f. Visual system histological sections stained with nuclear marker DAPI and phalloidin-Alexa488 for actin filaments from wildtype (a, c and e, n=5 for both stages) and *vsx*KO (b, d and f) animals at 6dpf (a, b, n=8), and 19hpf (c, e, d and f, n=6). No changes in eye size are observed but lamination in *vsx* mutants is severely affected at later stages compared to siblings (e, f). dpf: days post fertilization, ONL: outer nuclear layer, INL: inner nuclear layer, GCL: ganglion cell layer. Scale bar a-d: 100μm.

**VideoS1.**
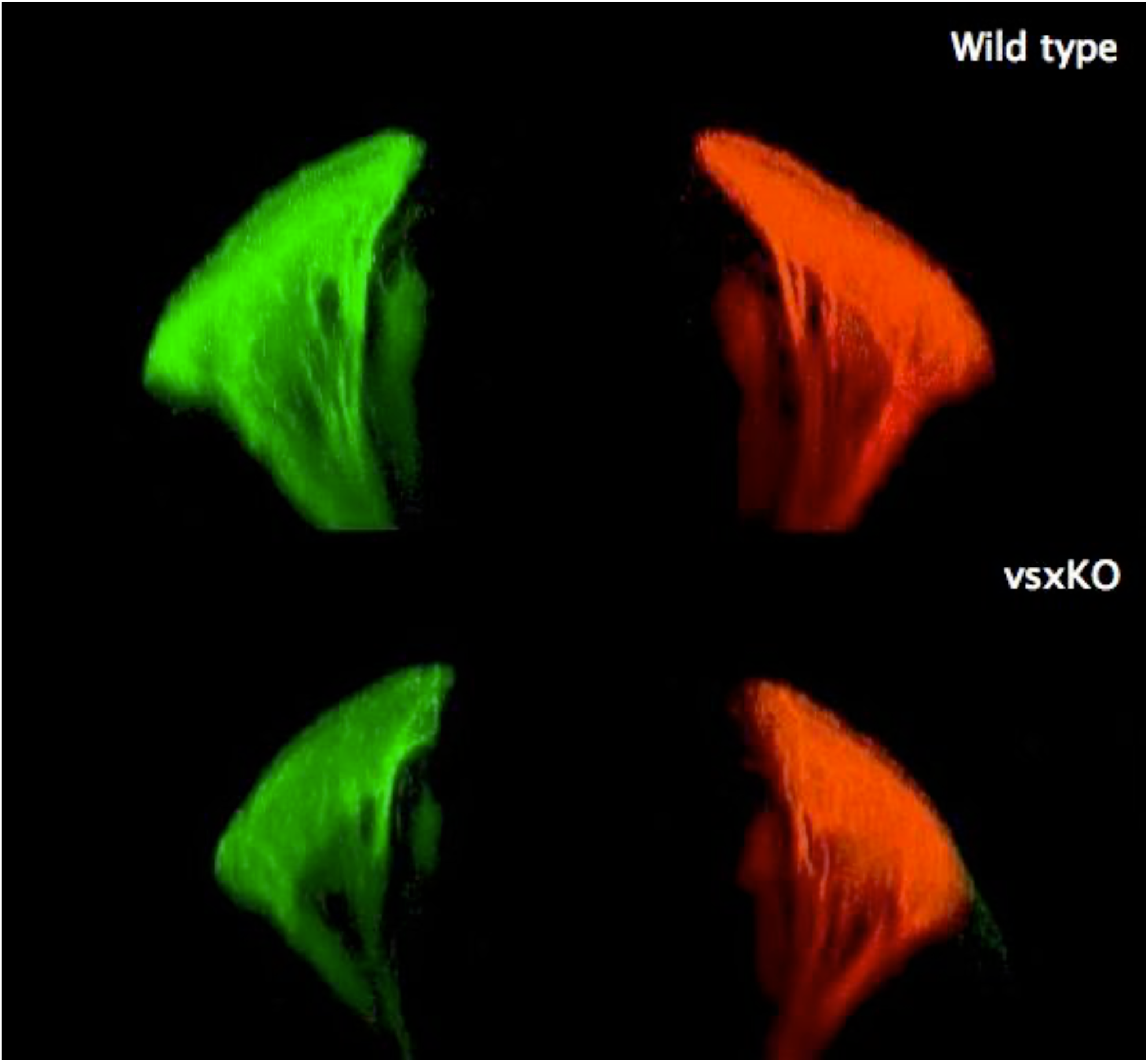
*vsx*KO larvae show normal GCL retinotectal projections. a, b. 3-D reconstructions of confocal stacks from zebrafish larval eyes injected with either DiO (green) or DiI (red) to label retinal ganglion cells and their projections to the optic tectum in wildtype (a, n=6) and *vsx*KO (b, n=8) at 6dpf. Note that *vsx*KO larvae show aparently normal retinotectal projections.

## METHODS

### Animal Experimentation and strains

All experiments performed in this work comply European Community standards for the use of animals in experimentation and were approved by ethical committees from Universidad Pablo de Olavide, Consejo Superior de Investigaciones Científicas (CSIC), the Andalusian government and Universidad Mayor. Zebrafish AB/Tübingen (AB/TU) and medaka iCab wild-type strains were staged, maintained and bred under standard conditions (Iwamatsu, 2004; Kimmel et al., 1995). Zebrafish Vsx mutants were maintained harbouring a single copy of *vsx1* (*vsx1*Δ245+/-, *vsx2*Δ73-/-) or *vsx2* (*vsx1*Δ245-/-;*vsx2*Δ73+/-), while medaka Vsx mutants were maintained in heterozygosis (*vsx1*Δ148+/-, *vsx2.1*Δ319+/-).

### Gene editing

Single guide RNAs (sgRNAs) targeting the DNA binding domains of *vsx1* and *vsx2* from zebrafish and *vsx1* and *vsx2.1* from medaka were designed using the CRISPRscan (Moreno-Mateos et al., 2015) and CCTop (Stemmer et al., 2015) design tools. Primers for sgRNA generation (see table1), were aligned by PCR to a universal CRISPR primer and the PCR product was further purified and used as template to sgRNA synthesis (Vejnar et al., 2016). To target individual *vsx* genes, a solution containing two sgRNAs (40 ng/μL each) and Cas9 protein (250 ng/μL) (Addgene; 47327) (Gagnon et al., 2014) were injected into one-cell-stage zebrafish and medaka embryos. Oligos used for screening of genomic DNA deletions flanking CRISPR target sites are detailed in table 1. Wild-type and mutant PCR products from F1 embryos were further analysed by standard sequencing to determine germline mutations (Stab Vida).

**Table 1.**
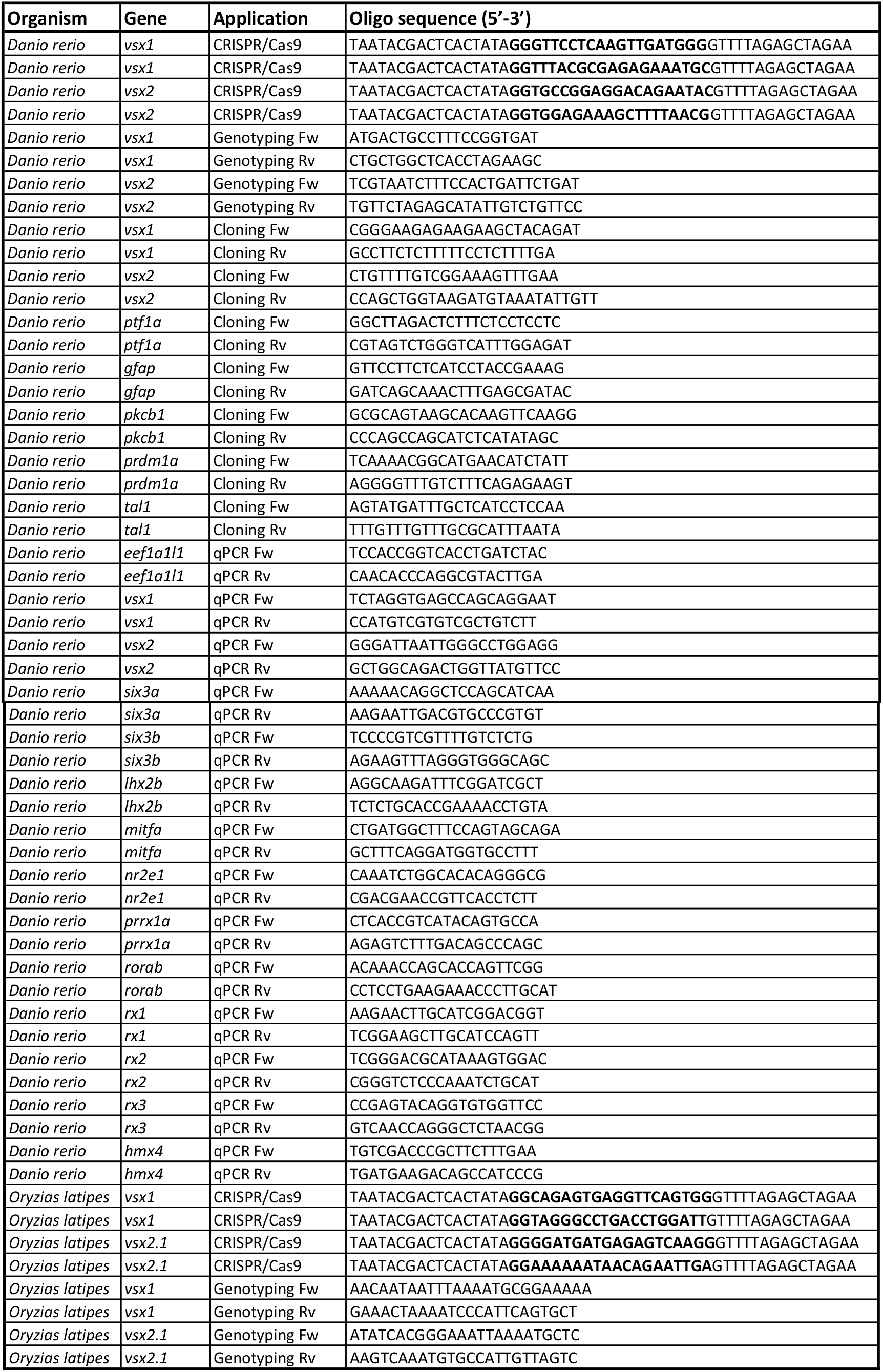
Nucleotide sequence of oligos used in this work. Organism, gene of interest, application and nucleotide sequence is described in each column. Note that the target site is bolded in CRISPR/Cas9 primers used for *vsx* disruption.

### Histology

Zebrafish and medaka samples from different developmental stages harbouring mutations in *vsx* genes were deeply anesthetized for 5-10 minutes with 160 mg/L of tricaine (ethyl 3-aminobenzoate methanesulfonate salt; MS-222; Merck) before dissecting their heads. Heads including both eyes were fixed in 4% w/v paraformaldehyde (PFA, Merck) in 0.1M phosphate buffer overnight at 4°C and the remaining tissue were kept for genotyping by conventional PCR. Wild-type and *vsx*KO sorted heads were then washed several times in 1X PBS, incubated in 30% sucrose-PBS overnight at 4°C, embedded in OCT (Tissue Tek) using cryomolds (Tissue Tek) and frozen in liquid nitrogen for short term storage at −80°C. Cryosectioning of samples was performed using a Leica CM1850 cryostat and 20μm thick transverse sections were collected in glass slides (Super Frost Ultra Plus, #11976299, Thermo Scientific) for Phalloidin (#A12379, Alexa fluor 488, Thermo Fisher Scientific) and 4’,6-Diamidine-2’-phenylindole dihydrochloride (DAPI, #10236276001, Merck) staining. Briefly, zebrafish and medaka eye transverse cryosections were dried at room temperature for ≥3h, and washed with filtered PBST (0.1% Triton in 1X PBS) 5 times for 5 minutes each wash. Then, slides were incubated with a solution containing 1/50 phalloidin alexa fluor 488 in PBST supplemented with 5% DMSO (Merck) and covered with parafilm (Bemis) in a dark humid chamber overnight at 4°C. After 30-60 minutes at room temperature, sections were incubated in a DAPI solution (1:1000 in PBST) and then washed 5 times for 5 minutes each wash with PBST. Slides were mounted with a drop of 15% glycerol in PBS and covered with 22×60mm coverslips. Mounted slides were kept in the dark and confocal images were captured immediately (≤24h) using a Leica SPE microscope to detect alexa 488 and DAPI signals from retina samples.

### Eye size and retina layer width measurements

Zebrafish embryos obtained from in-crosses of either *vsx1*Δ245+/-, *vsx2*Δ73-/- or *vsx1*Δ245-/-; *vsx2*Δ73+/- fish, were raised for 2 weeks under standard conditions (Kimmel et al., 1995). At this stage larvae were anesthetized, the antero-posterior length was measured (in millimetres) and a lateral image of the head region was obtained (Olympus SZX16 binocular scope connected to an Olympus DP71 camera). In parallel, a tip of the tail was collected using a scalpel to extract genomic DNA using Chelex resin (C7901, Sigma) for PCR screening. Head images (all taken at the same magnification) sorted by their genotype (either wild-type or *vsx*KO) were analysed using Fiji software to measure eye surface. Total eye area was divided by fish anteroposterior length for each animal to normalize eye size.

To measure retina INL and ONL layers width in zebrafish larvae, confocal images of eye cryostat sections from previously genotyped wild-type and *vsx*KO animals were taken using an immersion oil 40X objective (SPE, Leica). These images were then analysed using Fiji software to measure INL and ONL width (μm).

### Electroretinography (ERG)

ERG was recorded on 5 dpf larvae as previously described (Zang et al., 2015). 100 ms light stimuli delivered by HPX-2000 (Ocean Optics) were attenuated (log-4 to log0) by neutral density filters and given with an interval of 15 s. Full light intensity was measured by spectrometer (Ocean Optics, USB2000+) with spectrum shown in S1 (SpectraSuite, Ocean Optics). Electronic signals were amplified 1000 times by a pre-amplifier (P55 A.C. Preamplifier, Astro-Med. Inc, Grass Technology), digitized by DAQ Board (SCC-68, National Instruments) and recorded by self-written Labview program (National Instruments). Figures were prepared using Microsoft Excel 2016.

### Optokinetic response (OKR)

The OKR was recorded by the experiment setup as previously described (Mueller & Neuhauss, 2010). Briefly, 5dpf larvae were stimulated binocularly with sinusoidal gratings. To determine the contrast sensitivity, a spatial frequency of 20 cycles/360° and an angular velocity of 7.5 deg/s were used with varying contrast (5%, 10%, 20%, 40%, 70% and 100%). To study the spatial sensitivity, an angular velocity of 7.5/s and 70% of the maximum contrast was used with different spatial frequency (7, 14, 21,28,42,56 cycles/360°). To analyse the temporal sensitivity, maximum contrast and a spatial frequency of 20 cycles/360° were applied with increasing temporal frequency (5, 10, 15, 20, 25, 30 deg/s). Figures were presented by SPSS (Version 23.0. Armonk, NY: IBM Corp).

### Immunohistochemistry

Zebrafish wild-type and *vsx*KO retina sections from different developmental stages were analysed for the detection of apoptotic and mitotic cells using rabbit anti-active caspase-3 antibodies (BD Biosciences, 559565) and rabbit anti-phospho-histone H3 antibodies (Merck Millipore, 06-570), respectively. For the detection of cone and rod photoreceptors, zpr1 (ZIRC) and zpr3 (ZIRC) antibodies were used, respectively. Briefly, eye transverse cryosections were dried at room temperature for ≥3 h, washed 5 times for 5 minutes each with PBST, blocked for ≥1 h with 10% fetal bovine serum in PBST and incubated overnight in a humid chamber at 4°C with the corresponding primary antibody. All primary antibodies were diluted 1:500 in blocking solution. After several washes with PBST, a 1:500 dilution of the secondary antibody (Alexa Fluor 555 goat anti-rabbit or goat anti-mouse antibodies, Thermo Fisher, #A-21429 and #A-21422, respectively) was added for 2 h at room temperature. Following extensive washes with PBST, slides were mounted in 15% glycerol/PBS solution and sealed with 22×60mm coverslips. Immunofluorescence confocal images were taken using a Leica SPE confocal microscope.

### Fluorescent RNA *in situ* hybridization

Fluorescence *in situ* hybridization experiments were performed as previously described (Bogdanović et al., 2012). Fragments of the *vsx1*, *vsx2*, *ptf1a*, *prdm1a*, *gfap*, *pkcb1* and *tal1* genes were PCR amplified from zebrafish cDNA (SuperScript IV VILO Master Mix ThermoFisher Scientific, #11756050) using specific primers (table 1). For *vsx1* and *vsx2* genes, the deleted region of the coding sequence in *vsx*KO mutants was excluded from the amplified fragment. PCR products were cloned into StrataClone PCR Cloning vector (Agilent, #240205), linearized with XbaI restriction enzyme (Takara, #1093B) and transcribed with a DIG-labelling Kit (Roche, #11277073910) using T3 polymerase (Roche, #11031163001) to obtain digoxigenin-labelled antisense probes. Probes were used at a final concentration of 2ng/μl diluted in hybridization buffer (Thisse & Thisse, 2008).

### RNA-seq

Total RNA was extracted from 18 hpf zebrafish embryos’ heads using 1 ml TRIzol (Invitrogen, #15596026) following the manufacturer’s protocol. The trunk and tail of the embryos was used to extract genomic DNA using Chelex resin (C7901, Sigma) for PCR screening. Potential DNA contamination was eliminated by treating RNA samples with TURBO DNAse-free kit (Ambion, #AM1907). The concentration of the RNA samples was evaluated by Qubit spectrophotometer (Thermo Fisher). Libraries were prepared with TruSeq stranded mRNA kit (Illumina) and sequenced 2×125 bp on an Illumina Nextseq platform. We obtain at least 33 million reads per sample. Three biological replicates were used for each analysed condition. Reads were aligned to the danRer10 zebrafish genome assembly using hisat2 (Kim et al., 2015). Transcript abundance was estimated with Cufflinks v2.2.1. Differential gene expression analysis was performed using Cuffdiff v2.2.1, setting a corrected p-value < 0.05 as the cutoff for statistical significance of the differential expression. Multidimensional scaling analysis (MDS) was performed using the function MDSplot of the CummeRbund package in R 3.6.1.

### qPCR

cDNA retrotranscription and qPCR were performed as previously described (Vázquez-Marín et al., 2019). *eef1a1l1* gene was used as housekeeping normalization control. Primer sequences for amplified genes are detailed in table 1.

### ATAC-seq

Each ATAC-seq sample was obtained starting from a single 18 hpf zebrafish embryo’s head manually dissected, while the trunk and tail was kept for conventional PCR genotyping. Tagmentation and library amplification were performed using the FAST-ATAC protocol previously described (Corces et al., 2016). We obtained at least 70 M reads from the sequencing of each library. For data comparison, we used two biological replicates for each condition. Reads were aligned to the danRer10 zebrafish genome using Bowtie2 (Langmead & Salzberg, 2012) with −X 2000—no-mixed—no-unal parameters. PCR artifacts and duplicates were removed with the tool rmdup, available in the Samtools toolkit (Li et al., 2009). In order to detect the exact position where the transposase binds to the DNA, read start sites were offset by +4/-5 bp in the plus and minus strands. Read pairs that have an insert <130 bp were selected as nucleosome-free reads. Differential chromatin accessibility was calculated as reported (Magri et al., 2020). All chromatin regions reporting differential accessibility with an adjusted p-value < 0.05 were considered as DOCRs. All the DOCRS have been associated with genes using the online tool GREAT (McLean et al., 2010) with the option “basal plus extension”. Gene ontology enrichment analysis of the genes associated with DOCRs was performed with PANTHER (Mi et al., 2021).

### Labelling of retinotectal projections (DiI/DiO injections)

Following PCR genotyping, 6 dpf wild-type and *vsx* mutant larvae were fixed in 4% PFA overnight, washed in PBS and embedded in 1% low melting agarose (Sigma, #A9414) in PBS on an agarose filled Petri dish for injection. Each eye (between the lens and the retina) of the fish was injected either with 1% DiI (Invitrogen, #D275) or 1% DiO (Invitrogen, #D282) solutions in Chloroform (or dimethylformamide) with a pulled capillary glass mounted on a micromanipulator and under a stereomicroscope. A PV820 Pneumatic PicoPump (WPI) with the appropriate setting to deliver pressure to label the whole retina was used. Injected simples where washed in PBS, maintained overnight at 4°C and mounted on low melting agarose to image on a Zeiss LSM 710 confocal microscope. Z-stacks (0.5 μm x 0.5 μm x 1 μm) were collected to visualize the optic nerve reaching the tectum and 3D reconstructions were generated using Zen blue edition software (Zeiss).

### Statistical analysis

Quantitative data were evaluated using Prism 9.0 GraphPad software. Two-way ANOVA and a Tukey post hoc test was used to analyse ERG data, one-way ANOVA for OKR recordings and an unpaired *t* test for PH3, C3, eye area, retina layers’ width and trunk V2 neuron comparisons, were used. n values and significance levels are indicated in figure legends.

## Data availability

All datasets from this work are available in the Gene Expression Omnibus (GEO) repository (https://www.ncbi.nlm.nih.gov/geo/) under the following accession number: <u>GSE189739</u>. This Super series is composed of the following sub series: RNA-seq data are available under the GEO accession number <u>GSE189737</u> and ATAC-seq data are available under <u>GSE189738</u> number.

## Acknowledgments

We thank Marta Magri and Jorge Corbacho, for their scientific advice and Constanza Mounieres for technical assistance. We also thank the CABD Proteomics, Aquatic Vertebrates and Functional Genomics facilities for their excellent technical assistance. This work was supported by grants awarded to J.L. from ANID (FONDECYT Iniciación #11180727) and J.R.M.-M. from Junta de Andalucía (Reference PY20_00006), CSIC (Reference 2020AEP014), and Spanish Ministry of Science, Innovation and Universities: (References BFU2017-86339P, RED2018-102553-T, PID2020-112566GB-I00, and CEX2020-001088-M).

## REFERENCES

Amy Chen, C. M., & Cepko, C. L. (2000). Expression of Chx10 and Chx10-1 in the developing chicken retina. Mechanisms of Development, 90(2), 293–297. https://doi.org/10.1016/S0925-4773(99)00251-8

Bar-Yosef, U., Abuelaish, I., Harel, T., Hendler, N., Ofir, R., & Birk, O. S. (2004). CHX10 mutations cause non-syndromic microphthalmia/ anophthalmia in Arab and Jewish kindreds. Human Genetics, 115(4), 302–309. https://doi.org/10.1007/S00439-004-1154-2

Barabino, S. M. L., Spada, F., Cotelli, F., & Boncinelli, E. (1997). Inactivation of the zebrafish homologue of Chx10 by antisense oligonucleotides causes eye malformations similar to the ocular retardation phenotype. Mechanisms of Development, 63(2), 133–143. https://doi.org/10.1016/S0925-4773(97)00036-1

Bassett, E. A., & Wallace, V. A. (2012). Cell fate determination in the vertebrate retina. Trends in Neurosciences, 35(9), 565–573. https://doi.org/10.1016/J.TINS.2012.05.004

Bernardos, R. L., & Raymond, P. A. (2006). GFAP transgenic zebrafish. Gene Expression Patterns: GEP, 6(8), 1007–1013. https://doi.org/10.1016/J.MODGEP.2006.04.006

Bilotta, J., Saszik, S., & Sutherland, S. E. (2001). Rod contributions to the electroretinogram of the dark-adapted developing zebrafish. Developmental Dynamics: An Official Publication of the American Association of Anatomists, 222(4), 564–570. https://doi.org/10.1002/DVDY.1188

Bogdanović, O., Delfino-Machín, M., Nicolás-Pérez, M., Gavilán, M. P., Gago-Rodrigues, I., Fernández-Miñán, A., Lillo, C., Ríos, R. M., Wittbrodt, J., & Martínez-Morales, J. R. (2012). Numb/Numbl-Opo antagonism controls retinal epithelium morphogenesis by regulating integrin endocytosis. Developmental Cell, 23(4), 782–795. https://doi.org/10.1016/J.DEVCEL.2012.09.004

Brzezinski IV, J. A., Lamba, D. A., & Reh, T. A. (2010). Blimp1 controls photoreceptor versus bipolar cell fate choice during retinal development. Development, 137(4), 619–629. https://doi.org/10.1242/dev.043968

Buono, L., Corbacho, J., Naranjo, S., Almuedo-Castillo, M., Moreno-Marmol, T., de la Cerda, B., Sanbria-Reinoso, E., Polvillo, R., Díaz-Corrales, F. J., Bogdanovic, O., Bovolenta, P., & Martínez-Morales, J. R. (2021). Analysis of gene network bifurcation during optic cup morphogenesis in zebrafish. Nature Communications, 12(1). https://doi.org/10.1038/S41467-021-24169-7

Buono, L., & Martinez-Morales, J. R. (2020). Retina Development in Vertebrates: Systems Biology Approaches to Understanding Genetic Programs: On the Contribution of Next-Generation Sequencing Methods to the Characterization of the Regulatory Networks Controlling Vertebrate Eye Development. BioEssays, 42(4), 1–9. https://doi.org/10.1002/bies.201900187

Burmeister, M., Novak, J., Liang, M. Y., Basu, S., Ploder, L., Hawes, N. L., Vidgen, D., Hoover, F., Goldman, D., Kalnins, V. I., Roderick, T. H., Taylor, B. A., Hankin, M. H., & McInnes, R. R. (1996). Ocular retardation mouse caused by Chx10 homeobox null allele: impaired retinal progenitor proliferation and bipolar cell differentiation. Nature Genetics 1996 12:4, 12(4), 376–384. https://doi.org/10.1038/ng0496-376

Capowski, E. E., Wright, L. S., Liang, K., Phillips, M. J., Wallace, K., Petelinsek, A., Hagstrom, A., Pinilla, I., Borys, K., Lien, J., Min, J. H., Keles, S., Thomson, J. A., & Gamm, D. M. (2016). Regulation of WNT Signaling by VSX2 During Optic Vesicle Patterning in Human Induced Pluripotent Stem Cells. Stem Cells (Dayton, Ohio), 34(11), 2625–2634. https://doi.org/10.1002/STEM.2414

Chow, R. L., Snow, B., Novak, J., Looser, J., Freund, C., Vidgen, D., Ploder, L., & McInnes, R. R. (2001). Vsx1, a rapidly evolving paired-like homeobox gene expressed in cone bipolar cells. Mechanisms of Development, 109(2), 315–322. https://doi.org/10.1016/S0925-4773(01)00585-8

Chow, R. L., Volgyi, B., Szilard, R. K., Ng, D., McKerlie, C., Bloomfield, S. A., Birch, D. G., & McInnes, R. R. (2004). Control of late off-center cone bipolar cell differentiation and visual signaling by the homeobox gene Vsx1. Proceedings of the National Academy of Sciences of the United States of America, 101(6), 1754–1759. https://doi.org/10.1073/PNAS.0306520101

Clark, A. M., Yun, S., Veien, E. S., Wu, Y. Y., Chow, R. L., Dorsky, R. I., & Levine, E. M. (2008). Negative regulation of Vsx1 by its paralog Chx10/Vsx2 is conserved in the vertebrate retina. Brain Research, 1192, 99–113. https://doi.org/10.1016/J.BRAINRES.2007.06.007

Corces, M. R., Buenrostro, J. D., Wu, B., Greenside, P. G., Chan, S. M., Koenig, J. L., Snyder, M. P., Pritchard, J. K., Kundaje, A., Greenleaf, W. J., Majeti, R., & Chang, H. Y. (2016). Lineagespecific and single-cell chromatin accessibility charts human hematopoiesis and leukemia evolution. Nature Genetics, 48(10), 1193–1203. https://doi.org/10.1038/NG.3646

Crone, S. A., Quinlan, K. A., Zagoraiou, L., Droho, S., Restrepo, C. E., Lundfald, L., Endo, T., Setlak, J., Jessell, T. M., Kiehn, O., & Sharma, K. (2008). Genetic ablation of V2a ipsilateral interneurons disrupts left-right locomotor coordination in mammalian spinal cord. Neuron, 60(1), 70–83. https://doi.org/10.1016/J.NEURON.2008.08.009

Dorval, K. M., Bobechko, B. P., Ahmad, K. F., & Bremner, R. (2005). Transcriptional activity of the paired-like homeodomain proteins CHX10 and VSX1. The Journal of Biological Chemistry, 280(11), 10100–10108. https://doi.org/10.1074/JBC.M412676200

El-Brolosy, M. A., Kontarakis, Z., Rossi, A., Kuenne, C., Günther, S., Fukuda, N., Kikhi, K., Boezio, G. L. M., Takacs, C. M., Lai, S. L., Fukuda, R., Gerri, C., Giraldez, A. J., & Stainier, D. Y. R. (2019). Genetic compensation triggered by mutant mRNA degradation. Nature, 568(7751), 193–197. https://doi.org/10.1038/S41586-019-1064-Z

Erclik, T., Hartenstein, V., Lipshitz, H. D., & McInnes, R. R. (2008). Conserved role of the Vsx genes supports a monophyletic origin for bilaterian visual systems. Current Biology: CB, 18(17), 1278–1287. https://doi.org/10.1016/J.CUB.2008.07.076

Ferda Percin, E., Ploder, L. A., Yu, J. J., Arici, K., Jonathan Horsford, D., Rutherford, A., Bapat, B., Cox, D. W., Duncan, A. M. V., Kalnins, V. I., Kocak-Altintas, A., Sowden, J. C., Traboulsi, E., Sarfarazi, M., & McInnes, R. R. (2000). Human microphthalmia associated with mutations in the retinal homeobox gene CHX10. Nature Genetics, 25(4), 397–401. https://doi.org/10.1038/78071

Fleisch, V. C., & Neuhauss, S. C. F. (2006). Visual behavior in zebrafish. Zebrafish, 3(2), 191–201. https://doi.org/10.1089/ZEB.2006.3.191

Focareta, L., Sesso, S., & Cole, A. G. (2014). Characterization of homeobox genes reveals sophisticated regionalization of the central nervous system in the European cuttlefish Sepia officinalis. PloS One, 9(10). https://doi.org/10.1371/JOURNAL.PONE.0109627

Fuhrmann, S. (2010). Eye morphogenesis and patterning of the optic vesicle. In Current Topics in Developmental Biology (Vol. 93, Issue C). https://doi.org/10.1016/B978-0-12-385044-7.00003-5

Fujitani, Y., Fujitani, S., Luo, H., Qiu, F., Burlison, J., Long, Q., Kawaguchi, Y., Edlund, H., MacDonald, R. J., Furukawa, T., Fujikado, T., Magnuson, M. A., Xiang, M., & Wright, C. V. E. (2006). Ptf1a determines horizontal and amacrine cell fates during mouse retinal development. Development (Cambridge, England), 133(22), 4439–4450. https://doi.org/10.1242/DEV.02598

Gagnon, J. A., Valen, E., Thyme, S. B., Huang, P., Ahkmetova, L., Pauli, A., Montague, T. G., Zimmerman, S., Richter, C., & Schier, A. F. (2014). Efficient mutagenesis by Cas9 protein-mediated oligonucleotide insertion and large-scale assessment of single-guide RNAs. PLoS ONE, 9(5), 5–12. https://doi.org/10.1371/journal.pone.0098186

Gago-Rodrigues, I., Fernández-Miñán, A., Letelier, J., Naranjo, S., Tena, J. J., Gómez-Skarmeta, J. L., & Martinez-Morales, J. R. (2015). Analysis of opo cis-regulatory landscape uncovers Vsx2 requirement in early eye morphogenesis. Nature Communications, 6, 7054. https://doi.org/10.1038/ncomms8054

Goldman, D. (2014). Müller glial cell reprogramming and retina regeneration. Nature Reviews. Neuroscience, 15(7), 431–442. https://doi.org/10.1038/NRN3723

Goodson, N. B., Kaufma, M. A., Par, K. U., & Brzezinski, J. A. (2020). Simultaneous deletion of Prdm1 and Vsx2 enhancers in the retina alters photoreceptor and bipolar cell fate specification, yet differs from deleting both genes. Development (Cambridge), 147(13). https://doi.org/10.1242/dev.190272

Green, E. S., Stubbs, J. L., & Levine, E. M. (2003). Genetic rescue of cell number in a mouse model of microphthalmia: interactions between Chx10 and G1-phase cell cycle regulators. Development (Cambridge, England), 130(3), 539–552. https://doi.org/10.1242/DEV.00275

Gregory-Evans, C Y, Williams, M. J., Halford, S., & Gregory-Evans, K. (2004). Ocular coloboma: a reassessment in the age of molecular neuroscience. J Med Genet, 41, 881–891. https://doi.org/10.1136/jmg.2004.025494

Gregory-Evans, Cheryl Y., Wallace, V. A., & Gregory-Evans, K. (2013). Gene networks: Dissecting pathways in retinal development and disease. Progress in Retinal and Eye Research, 33(1), 40–66. https://doi.org/10.1016/J.PRETEYERES.2012.10.003

Heavner, W., & Pevny, L. (2012). Eye development and retinogenesis. Cold Spring Harbor Perspectives in Biology, 4(12). https://doi.org/10.1101/cshperspect.a008391

Héon, E., Greenberg, A., Kopp, K. K., Rootman, D., Vincent, A. L., Billingsley, G., Priston, M., Dorval, K. M., Chow, R. L., McInnes, R. R., Heathcote, G., Westall, C., Sutphin, J. E., Semina, E., Bremner, R., & Stone, E. M. (2002). VSX1: a gene for posterior polymorphous dystrophy and keratoconus. Human Molecular Genetics, 11(9), 1029–1036. https://doi.org/10.1093/HMG/11.9.1029

Horsford, D. J., Nguyen, M. T. T., Sellar, G. C., Kothary, R., Arnheiter, H., & McInnes, R. R. (2005). Chx10 repression of Mitf is required for the maintenance of mammalian neuroretinal identity. Development, 132(1), 177–187. https://doi.org/10.1242/dev.01571

Iwamatsu, T. (2004). Stages of normal development in the medaka Oryzias latipes. Mechanisms of Development, 121(7–8), 605–618. https://doi.org/10.1016/j.mod.2004.03.012

Jusuf, P. R., & Harris, W. A. (2009). Ptf1a is expressed transiently in all types of amacrine cells in the embryonic zebrafish retina. Neural Development, 4(1). https://doi.org/10.1186/1749-8104-4-34

Katoh, K., Omori, Y., Onishi, A., Sato, S., Kondo, M., & Furukawa, T. (2010). Blimp1 suppresses Chx10 expression in differentiating retinal photoreceptor precursors to ensure proper photoreceptor development. Journal of Neuroscience, 30(19), 6515–6526. https://doi.org/10.1523/JNEUROSCI.0771-10.2010

Kim, D., Langmead, B., & Salzberg, S. L. (2015). HISAT: a fast spliced aligner with low memory requirements. Nature Methods, 12(4), 357–360. https://doi.org/10.1038/NMETH.3317

Kim, D. S., Matsuda, T., & Cepko, C. L. (2008). A core paired-type and POU homeodomain-containing transcription factor program drives retinal bipolar cell gene expression. The Journal of Neuroscience: The Official Journal of the Society for Neuroscience, 28(31), 7748–7764. https://doi.org/10.1523/JNEUROSCI.0397-08.2008

Kimmel, C. B., Ballard, W. W., Kimmel, S. R., Ullmann, B., & Schilling, T. F. (1995). Stages of embryonic development of the zebrafish. Developmental Dynamics, 203(3), 253–310. https://doi.org/10.1002/aja.1002030302

Kimura, Y., Satou, C., Fujioka, S., Shoji, W., Umeda, K., Ishizuka, T., Yawo, H., & Higashijima, S. I. (2013). Hindbrain V2a neurons in the excitation of spinal locomotor circuits during zebrafish swimming. Current Biology: CB, 23(10), 843–849.https://doi.org/10.1016/J.CUB.2013.03.066

Kimura, Y., Satou, C., & Higashijima, S. I. (2008). V2a and V2b neurons are generated by the final divisions of pair-producing progenitors in the zebrafish spinal cord. Development, 135(18), 3001–3005. https://doi.org/10.1242/dev.024802

Kok, F. O., Shin, M., Ni, C. W., Gupta, A., Grosse, A. S., vanImpel, A., Kirchmaier, B. C., Peterson-Maduro, J., Kourkoulis, G., Male, I., DeSantis, D. F., Sheppard-Tindell, S., Ebarasi, L., Betsholtz, C., Schulte-Merker, S., Wolfe, S. A., & Lawson, N. D. (2015). Reverse genetic screening reveals poor correlation between morpholino-induced and mutant phenotypes in zebrafish. Developmental Cell, 32(1), 97–108. https://doi.org/10.1016/j.devcel.2014.11.018

Langmead, B., & Salzberg, S. L. (2012). Fast gapped-read alignment with Bowtie 2. Nature Methods, 9(4), 357–359. https://doi.org/10.1038/NMETH.1923

Li, H., Handsaker, B., Wysoker, A., Fennell, T., Ruan, J., Homer, N., Marth, G., Abecasis, G., & Durbin, R. (2009). The Sequence Alignment/Map format and SAMtools. Bioinformatics (Oxford, England), 25(16), 2078–2079. https://doi.org/10.1093/BIOINFORMATICS/BTP352

Liu, I. S. C., Chen, J. de, Ploder, L., Vidgen, D., van der Kooy, D., Kalnins, V. I., & Mclnnes, R. R. (1994). Developmental expression of a novel murine homeobox gene (Chx10): evidence for roles in determination of the neuroretina and inner nuclear layer. Neuron, 13(2), 377–393. https://doi.org/10.1016/0896-6273(94)90354-9

Livne-Bar, I., Pacal, M., Cheung, M. C., Hankin, M., Trogadis, J., Chen, D., Dorval, K. M., & Bremner, R. (2006). Chx10 is required to block photoreceptor differentiation but is dispensable for progenitor proliferation in the postnatal retina. Proceedings of the National Academy of Sciences of the United States of America, 103(13), 4988–4993. https://doi.org/10.1073/pnas.0600083103

Magri, M. S., Jiménez-Gancedo, S., Bertrand, S., Madgwick, A., Escrivà, H., Lemaire, P., & Gómez-Skarmeta, J. L. (2020). Assaying Chromatin Accessibility Using ATAC-Seq in Invertebrate Chordate Embryos. Frontiers in Cell and Developmental Biology, 7. https://doi.org/10.3389/FCELL.2019.00372

Martinez-Morales, J. R. (2016). Vertebrate eye gene regulatory networks. Organogenetic Gene Networks: Genetic Control of Organ Formation, 259–274. https://doi.org/10.1007/978-3-319-42767-6_9

Matías-Pérez, D., García-Montaño, L. A., Cruz-Aguilar, M., García-Montalvo, I. A., Nava-Valdéz, J., Barragán-Arevalo, T., Villanueva-Mendoza, C., Villarroel, C. E., Guadarrama-Vallejo, C., la Cruz, R. V. de, Chacón-Camacho, O., & Zenteno, J. C. (2018). Identification of novel pathogenic variants and novel gene-phenotype correlations in Mexican subjects with microphthalmia and/or anophthalmia by next-generation sequencing. Journal of Human Genetics, 63(11), 1169–1180. https://doi.org/10.1038/S10038-018-0504-1

McLean, C. Y., Bristor, D., Hiller, M., Clarke, S. L., Schaar, B. T., Lowe, C. B., Wenger, A. M., & Bejerano, G. (2010). GREAT improves functional interpretation of cis-regulatory regions. Nature Biotechnology, 28(5), 495–501. https://doi.org/10.1038/NBT.1630

Meyer, A., & Schartl, M. (1999). Gene and genome duplications in vertebrates: the one-to-four (-to-eight in fish) rule and the evolution of novel gene functions. Current Opinion in Cell Biology, 11(6), 699–704. https://doi.org/10.1016/S0955-0674(99)00039-3

Mi, H., Ebert, D., Muruganujan, A., Mills, C., Albou, L. P., Mushayamaha, T., & Thomas, P. D. (2021). PANTHER version 16: a revised family classification, tree-based classification tool, enhancer regions and extensive API. Nucleic Acids Research, 49(D1), D394–D403. https://doi.org/10.1093/NAR/GKAA1106

Mintz-Hittner, H. A., Semina, E. V., Frishman, L. J., Prager, T. C., & Murray, J. C. (2004). VSX1 (RINX) mutation with craniofacial anomalies, empty sella, corneal endothelial changes, and abnormal retinal and auditory bipolar cells. Ophthalmology, 111(4), 828–836. https://doi.org/10.1016/J.OPHTHA.2003.07.006

Moreno-Mateos, M. A., Vejnar, C. E., Beaudoin, J. D., Fernandez, J. P., Mis, E. K., Khokha, M. K., & Giraldez, A. J. (2015). CRISPRscan: Designing highly efficient sgRNAs for CRISPR-Cas9 targeting in vivo. Nature Methods, 12(10), 982–988. https://doi.org/10.1038/nmeth.3543

Mueller, K. P., & Neuhauss, S. C. F. (2010). Quantitative measurements of the optokinetic response in adult fish. Journal of Neuroscience Methods, 186(1), 29–34. https://doi.org/10.1016/J.JNEUMETH.2009.10.020

Nguyen, M. T. T., & Arnheiter, H. (2000). Signaling and transcriptional regulation in early mammalian eye development: A link between FGF and MITF. Development, 127(16), 3581–3591. https://doi.org/10.1242/dev.127.16.3581

Norrie, J. L., Lupo, M. S., Xu, B., Al Diri, I., Valentine, M., Putnam, D., Griffiths, L., Zhang, J., Johnson, D., Easton, J., Shao, Y., Honnell, V., Frase, S., Miller, S., Stewart, V., Zhou, X., Chen, X., & Dyer, M. A. (2019). Nucleome Dynamics during Retinal Development. Neuron, 104(3), 512–528.e11. https://doi.org/10.1016/J.NEURON.2019.08.002

Ohtoshi, A., Wang, S. W., Maeda, H., Saszik, S. M., Frishman, L. J., Klein, W. H., & Behringer, R. R. (2004). Regulation of retinal cone bipolar cell differentiation and photopic vision by the CVC homeobox gene Vsx1. Current Biology: CB, 14(6), 530–536. https://doi.org/10.1016/J.CUB.2004.02.027

Osipov, V. V., & Vakhrusheva, M. P. (1983). Variation in the expressivity of the ocular retardation gene in mice. TSitologiia i Genetika, 17(4), 39–43. https://pubmed.ncbi.nlm.nih.gov/6623630/

Passini, M. A., Levine, E. M., Canger, A. K., Raymond, P. A., & Schechter, N. (1997). Vsx-1 and vsx-2: Differential expression of two paired-like homeobox genes during zebrafish and goldfish retinogenesis. Journal of Comparative Neurology, 388(3), 495–505. https://doi.org/10.1002/(SICI)1096-9861(19971124)388:3<495::AID-CNE11>3.0.CO;2-L

Reinhardt, R., Centanin, L., Tavhelidse, T., Inoue, D., Wittbrodt, B., Concordet, J., Martinez-Morales, J. R., & Wittbrodt, J. (2015). Sox2, Tlx, Gli3, and Her9 converge on Rx2 to define retinal stem cells in vivo. The EMBO Journal, 34(11), 1572–1588. https://doi.org/10.15252/EMBJ.201490706

Rinner, O., Rick, J. M., & Neuhauss, S. C. F. (2005). Contrast sensitivity, spatial and temporal tuning of the larval zebrafish optokinetic response. Investigative Ophthalmology & Visual Science, 46(1), 137–142. https://doi.org/10.1167/IOVS.04-0682

Rossi, A., Kontarakis, Z., Gerri, C., Nolte, H., Hölper, S., Krüger, M., & Stainier, D. Y. R. (2015). Genetic compensation induced by deleterious mutations but not gene knockdowns. Nature, 524(7564), 230–233. https://doi.org/10.1038/nature14580

Rowan, S., & Cepko, C. L. (2005). A POU factor binding site upstream of the Chx10 homeobox gene is required for Chx10 expression in subsets of retinal progenitor cells and bipolar cells. Developmental Biology, 281(2), 240–255. https://doi.org/10.1016/J.YDBIO.2005.02.023

Rowan, S., Chen, C. M. A., Young, T. L., Fisher, D. E., & Cepko, C. L. (2004). Transdifferentiation of the retina into pigmented cells in ocular retardation mice defines a new function of the homeodomain gene Chx10. Development, 131(20), 5139–5152. https://doi.org/10.1242/dev.01300

Satow, T., Bae, S. K., Inoue, T., Inoue, C., Miyoshi, G., Tomita, K., Bessho, Y., Hashimoto, N., & Kageyama, R. (2001). The basic helix-loop-helix gene hesr2 promotes gliogenesis in mouse retina. The Journal of Neuroscience: The Official Journal of the Society for Neuroscience, 21(4), 1265–1273. https://doi.org/10.1523/JNEUROSCI.21-04-01265.2001

Seth, A., Culverwell, J., Walkowicz, M., Toro, S., Rick, J. M., Neuhauss, S. C. F., Varga, Z. M., & Karlstrom, R. O. (2006). belladonna/(Ihx2) is required for neural patterning and midline axon guidance in the zebrafish forebrain. Development (Cambridge, England), 133(4), 725–735. https://doi.org/10.1242/DEV.02244

Shi, Z., Trenholm, S., Zhu, M., Buddingh, S., Star, E. N., Awatramani, G. B., & Chow, R. L. (2011). Vsx1 regulates terminal differentiation of type 7 ON bipolar cells. The Journal of Neuroscience: The Official Journal of the Society for Neuroscience, 31(37), 13118–13127. https://doi.org/10.1523/JNEUROSCI.2331-11.2011

Stemmer, M., Thumberger, T., Del Sol Keyer, M., Wittbrodt, J., & Mateo, J. L. (2015). CCTop: An Intuitive, Flexible and Reliable CRISPR/Cas9 Target Prediction Tool. PloS One, 10(4). https://doi.org/10.1371/JOURNAL.PONE.0124633

Thisse, C., & Thisse, B. (2008). High-resolution in situ hybridization to whole-mount zebrafish embryos. Nature Protocols, 3(1), 59–69. https://doi.org/10.1038/NPROT.2007.514

Truslove, G. M. (1962). A gene causing ocular retardation in the mouse. Journal of Embryology and Experimental Morphology, 10, 652–660. https://doi.org/10.1242/dev.10.4.652

Vázquez-Marín, J., Gutiérrez-Triana, J. A., Almuedo-Castillo, M., Buono, L., Gómez-Skarmeta, J. L., Mateo, J. L., Wittbrodt, J., & Martínez-Morales, J. R. (2019). yap1b, a divergent Yap/Taz family member, cooperates with yap1 in survival and morphogenesis via common transcriptional targets. Development (Cambridge, England), 146(13). https://doi.org/10.1242/DEV.173286

Vejnar, C. E., Moreno-Mateos, M. A., Cifuentes, D., Bazzini, A. A., & Giraldez, A. J. (2016). Optimized CRISPR-Cas9 System for Genome Editing in Zebrafish. Cold Spring Harbor Protocols, 2016(10), 856–870. https://doi.org/10.1101/PDB.PROT086850

Vitorino, M., Jusuf, P. R., Maurus, D., Kimura, Y., Higashijima, S. I., & Harris, W. A. (2009). Vsx2 in the zebrafish retina: Restricted lineages through derepression. Neural Development, 4(1). https://doi.org/10.1186/1749-8104-4-14

Wang, S., Sengel, C., Emerson, M. M., & Cepko, C. L. (2014). A gene regulatory network controls the binary fate decision of rod and bipolar cells in the vertebrate retina. Developmental Cell, 30(5), 513–527. https://doi.org/10.1016/J.DEVCEL.2014.07.018

Wilm, T. P., & Solnica-Krezel, L. (2005). Essential roles of a zebrafish prdm1/blimp1 homolog in embryo patterning and organogenesis. Development (Cambridge, England), 132(2), 393–404. https://doi.org/10.1242/DEV.01572

Zang, J., Keim, J., Kastenhuber, E., Gesemann, M., & Neuhauss, S. C. F. (2015). Recoverin depletion accelerates cone photoresponse recovery. Open Biology, 5(8). https://doi.org/10.1098/RSOB.150086

